# WNK1 regulates uterine homeostasis and its ability to support pregnancy

**DOI:** 10.1101/2020.03.27.012039

**Authors:** Ru-pin Alicia Chi, Tianyuan Wang, Chou-Long Huang, San-pin Wu, Steven Young, John Lydon, Francesco DeMayo

## Abstract

WNK1 is an atypical kinase protein ubiquitously expressed in humans and mice. A mutation in its encoding gene causes hypertension in humans which is associated with abnormal ion homeostasis. Our earlier findings demonstrated that WNK1 is critical for *in vitro* decidualization in human endometrial stromal cells – pointing towards an unrecognized role of WNK1 in female reproduction. Here, we employed a mouse model with conditional WNK1 ablation from the female reproductive tract to define its *in vivo* role in uterine biology. Loss of WNK1 altered uterine morphology, causing endometrial epithelial hyperplasia, adenomyosis and a delay in embryo implantation, ultimately resulting in compromised fertility. Combining transcriptomic, proteomic and interactomic analyses revealed a novel regulatory pathway whereby WNK1 represses AKT phosphorylation through the phosphatase PP2A in endometrial cells from both humans and mice. We show that WNK1 interacts with PPP2R1A, an isoform of the PP2A scaffold subunit. This interaction stabilizes the PP2A complex, which then dephosphorylates AKT. Therefore, loss of WNK1 reduced PP2A activity, causing AKT hypersignaling. Using FOXO1 as a readout of AKT activity, we demonstrate that there was escalated FOXO1 phosphorylation and nuclear exclusion, leading to a disruption in the regulation of genes that are crucial for embryo implantation.

## Introduction

The ability of the uterus to support and maintain the development of an embryo is critically dependent upon the process of implantation, the aberrant occurrence of which causes a ripple effect leading to pregnancy complications and miscarriage (1). Embryo implantation occurs during the “window of receptivity” and requires a fully prepared and receptive uterus. Investigation of uterine function mediators during receptivity identified <u>w</u>ith <u>n</u>o lysine(<u>k</u>) kinase (K) 1 (WNK1) as functional in the uterus acting downstream of EGFR, whose inhibition impaired decidualization (2). Functional analysis of human endometrial stromal cells *in vitro* demonstrated that WNK1 is needed for proliferation, migration and differentiation (3). Collectively, these findings indicate a previously unrecognized function of WNK1 in the female reproductive tract and led us to hypothesize that WNK1 is a mediator of uterine function.

WNK1 belongs to a family of serine/threonine protein kinases (4, 5), with its name derived from the unusual placement of the catalytic lysine in subdomain I (6). To date, Wnk1’s function is the most extensively explored in the kidney and the nervous system due to the link between its mutation and familial hypertension and autonomic neuropathy (7-9). In the renal system, WNK1 controls ion homeostasis through diverse mechanisms including activation of the SGK1/epithelial sodium channel pathway (10), regulating the potassium channel Kir1.1 cell surface localization (11), as well as controlling the activity of Na-K-Cl cotransporter through phosphorylating OSR1(OXSR1) and SPAK (12, 13). Interestingly, WNK1’s regulatory function on OSR1 is critical for cardiovascular development, thereby contributing to embryonic lethality when WNK1 is ablated from the endothelium (14, 15). These findings suggest that although WNK1 exhibits organ-specific physiological functions, the underlying cellular components regulated by WNK1 may share similarity between the different tissues.

Despite its ubiquitous expression pattern, WNK1’s role in organs other than those described above remain unexplored. Given the role of WNK1 in regulating uterine stromal cell biology *in vitro*, we hypothesized that WNK1 is essential in regulating uterine functions. To test this idea, we established a conditional uterine WNK1 ablated mouse model, which resulted in severely compromised fertility. We demonstrate that WNK1 is critical in maintaining uterine morphology, regulating epithelial proliferation and permitting appropriate embryo implantation. Transcriptomic and proteomic analyses identified deregulation of the AKT signaling pathway underlying the observed phenotypes. Using cultured human endometrial cells, we conducted a series of functional analyses to tease out the signaling hierarchy involving WNK1, PP2A, AKT and FOXO1.

## RESULTS

### WNK1 is expressed in the uterus during early pregnancy in both humans and mice

WNK1 expression was examined by immunohistochemistry in human endometrium during the proliferative and mid-secretory phases as well as in the peri-implantation uterus of mice. In humans, WNK1 is expressed in both the epithelial and stromal cells during the proliferative and mid-secretory phases (Fig. 1.A). Similarly in mice, WNK1 is expressed during and after implantation on gestation days (GDs) 4.5 and 5.5 (Fig. 1.B). These findings support the *in vivo* involvement of WNK1 in regulating functions of the female reproductive tract.

**Figure 1.**
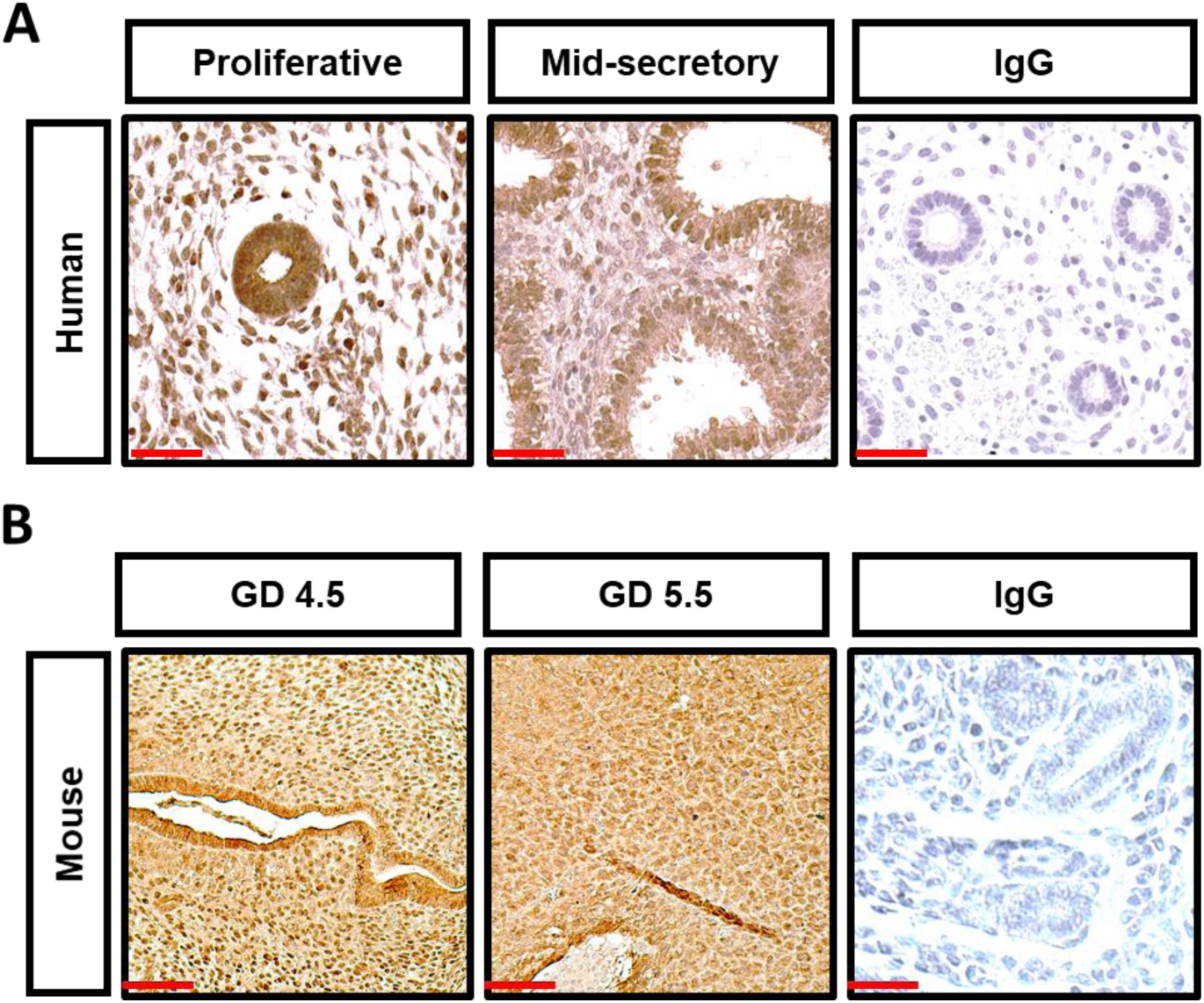
WNK1 is expressed in the uterus during the window of implantation in both humans and mice. (A and B) Immunohistochemical staining of WNK1 in proliferative and mid-secretory phased endometrial tissues from fertile women (A), and during receptive gestation day (GD) 4.5 and post implantation GD 5.5 in the uterus of wild-type mice (B). IgG served as negative controls, scale bars = 50 µm.

### WNK1 ablation altered uterine morphology and microenvironment

To examine WNK1’s function in the female reproductive tract, we established a mouse model with WNK1 ablation in progesterone receptor (PGR) expressing cells, which comprised all cellular compartments of the uterus. The conditional *Wnk1* allele mice (Wnk1^f/f^) were crossed to the PGR^Cre^ mice (14, 16), and confirmed that in the PGR^Cre/+^;Wnk1^f/f^ offspring (Wnk1^d/d^), Cre mediated excision led to the removal of exon 2 (Fig. S1.A and S1.B). Analysis of protein expression in uterine tissue via western blot confirmed reduced WNK1 protein levels in the Wnk1^d/d^ mice (Fig. S1.C), indicating that exon 2 excision led to reduced total protein levels.

We employed a tissue clearing technique to visualize uterine morphology in 3D during the window of receptivity (Fig. 2.A) (17). WNK1 deficiency substantially impacted uterine morphology, exhibiting increased number of, as well as altered structure of the endometrial glands (Fig. 2.A). Among the abnormalities observed in the Wnk1^d/d^ uteri was the failure of gland reorientation surrounding the embryo (18, 19). This is seen in the Wnk1^f/f^ uteri where glands near the embryo exhibited a elongated structure while glands away from the embryo remained tortuous and intertwined. In the Wnk1d/d uteri, the glands appears equally tortuous irrespective of distance from embryo (Fig. 2.A). Examination of uterine cross sections from older mice (26 and 50 weeks) further demonstrated invasion of glands into the myometrium, suggesting that WNK1 ablation caused adenomyosis (Fig. 2.B). This was supported by the elevated expression of *Moesin* (*Msn*) in the Wnk1^d/d^ uteri, a biomarker for adenomyosis in humans (Fig. 2.C) (20). Quantification of gland number and *Foxa2* gene expression showed significant elevation in the Wnk1^d/d^ uteri (Fig. 2.D and E), confirming the substantial increase in glandular tissues. To examine whether the increased glands were a result of increased proliferation in the uterus, we examined the expression of two mitotic markers - cyclin D1 (CCND1) and phosphorylated histone H3 (H3S10p). Elevated levels of both proteins in the glandular epithelial cells of the Wnk1^d/d^ uteri was observed (Fig. 2.F). In addition to the increased CCND1 and H3S10p in the glandular epithelium, higher expression of both proteins was also observed in the luminal epithelium of Wnk1^d/d^ uteri demonstrating that WNK1 ablation induced epithelial hyperplasia was not restricted to the glands. Moreover, we observed increased extracellular matrix deposition especially surrounding the glands in the Wnk1^d/d^ uteri, as shown by Masson’s trichrome staining (Fig. 2.G). These results suggest that the adenomyotic phenotype could be associated with increased epithelial proliferation as well as excessive extracellular matrix deposition (21, 22).

**Figure 2.**
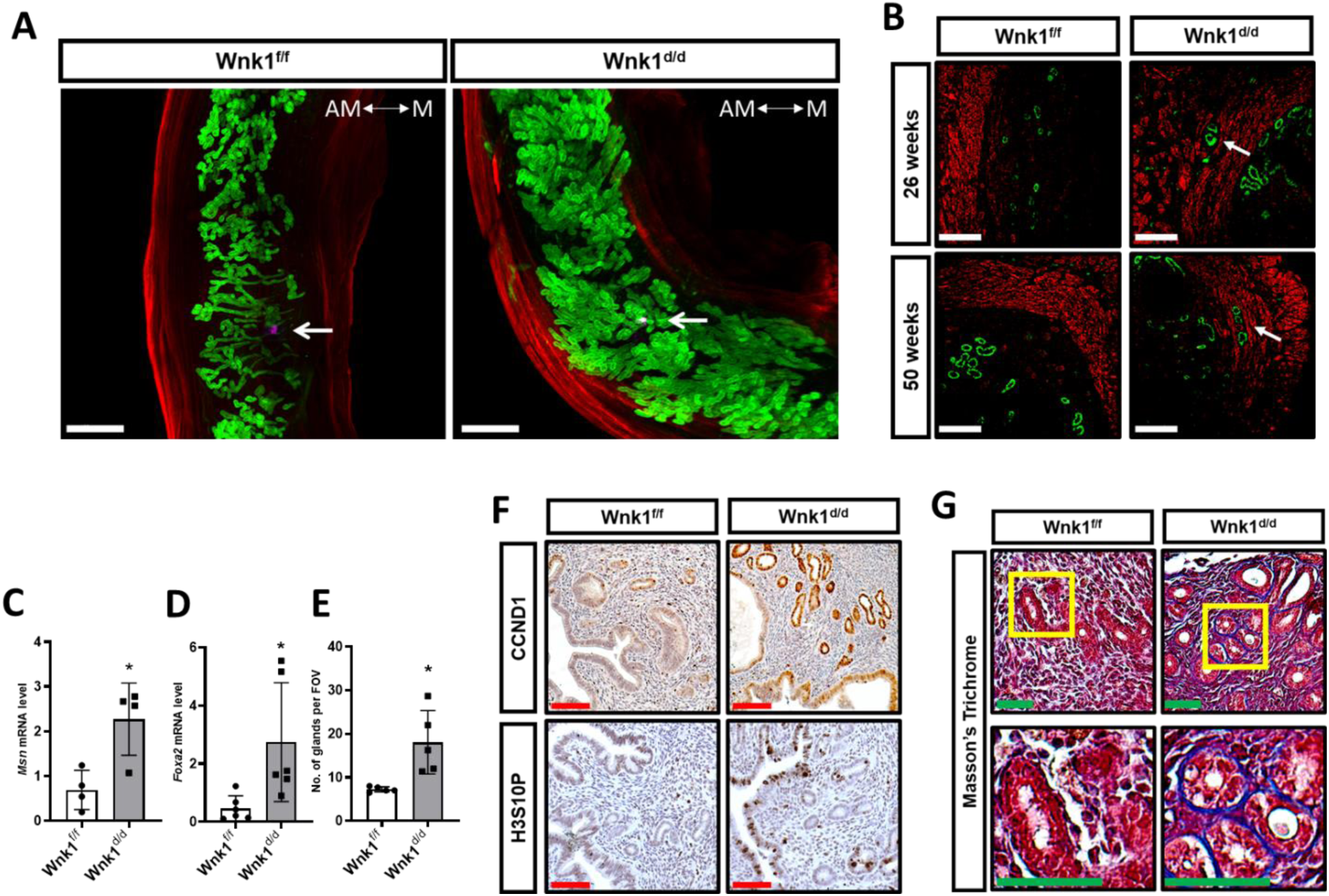
WNK1 ablation altered uterine morphology and microenvironment. (A) Three-dimensional reconstruction of Wnk1^f/f^ and Wnk1^d/d^ uteri on GD 4.5 using tissue clearing and confocal microscopy. The glands, myometrium and embryo were marked by FOXA2 (green), ACTA2 (red) and OCT4 (purple), respectively. Images were captured by tile-scanning and Z-stacking, and reassembled *in silico* using Imaris software. White arrow indicates position of the embryo. Scale bars = 500 µm. The antimesometrial (AM) and mesometrial (M) sides of the tissue are indicated. (B) Immunofluorescence of uterine cross section showing glands (FOXA2, green) and myometrium (ACTA2, red) from Wnk1^f/f^ and Wnk1^d/d^ uteri. White arrows indicate gland extension into myometrium. Scale bars = 50 µm. (C) Adenomyosis biomarker *Msn* mRNA expression as determined by qRT-PCR (n = 4). (D) Quantification of *Foxa2* mRNA expression as determined by qRT-PCR (n = 6), and (E) Quantification of number of glands per cross section for Wnk1^f/f^ and Wnk1^d/d^ mice (n = 6). (F) Expression of mitotic markers CCND1 and H3S10P in the uterus of 26-week-old Wnk1^f/f^ and Wnk1^d/d^ mice, scale bars = 100 µm. (G) Masson’s trichrome staining of uteri cross section from 26 and 50 week old Wnk1^f/f^ and WNk1^d/d^ mice, scale bars = 100 µm. Yellow boxes indicate region shown at higher magnification in lower panels. All quantitative results shown are average ± SD, * *p* < 0.05.

### Uterine loss of WNK1 impaired implantation

A six month breeding trial was conducted to determine the impact of WNK1 ablation on female fertility. Of the 8 control mice, 7 were able to complete the breeding trial, with 1 found dead midtrial. Necropsy showed neither pregnancy nor abnormality associated with the reproductive tract in this one mouse, indicating that the cause of death was not related to abnormal uterine function. The 7 mice produced 31 litters totaling 245 pups, which was equivalent to 4.4 litters and 35 pups per mouse during the 6 months (Fig. 3.A). In contrast, only 4 of the 8 Wnk1^d/d^ mice initiated in the trial were able to complete the trial. This was due to 4 females succumbing to complications during pregnancy or delivery. Of those, 2 had been found dead near term each carrying 2 pups, 1 was in dystocia and had to undergo euthanization, from which 5 pups were recovered. One was found ill and necropsy showed utero-abdominal fistula. Of the 4 females that completed the trial, 10 litters and 18 pups were produced, which averaged to 2.5 litters and 4.5 pups per mouse (Fig. 3.A). The average litter size was also significantly smaller, averaging 3.3 pups per litter in the Wnk1^d/d^ mice, compared to 7.7 in the control Wnk1^f/f^ mice (Fig. 3.B). While the Wnk1^f/f^ mice bred consistently, producing the last litters in the twentieth week of the trial, most of the Wnk1^d/d^ mice stopped breeding after 3 litters less than 15 weeks into the trial, with only one mouse producing after the twentieth week (Fig. 3.C). Taken together, these results illustrated compromised ability to support pregnancy and premature sterility with uterine loss of WNK1.

**Figure 3.**
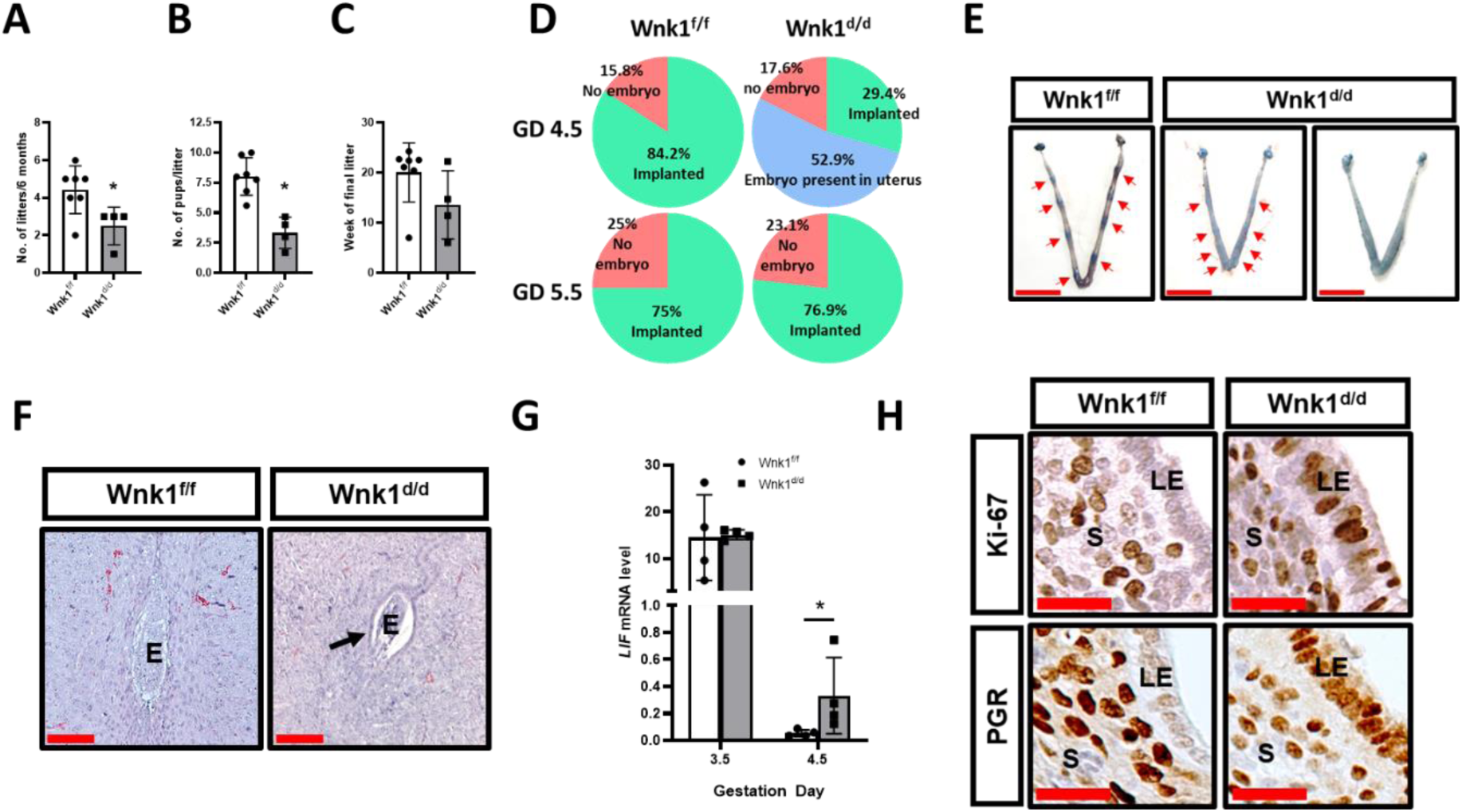
Uterine WNK1 ablation compromised fertility and impaired implantation in mice. (A -C) Results from a 6-month breeding trial where the Wnk1^f/f^ (n = 7) and Wnk1^d/d^ (n = 4) mice were housed with wildtype males, showing average number of litters produced, (B) average number of pups per litter produced and (C) time of last delivery. Results shown are mean ± SD, * *p* < 0.05. (D) Percentage of mated Wnk1^f/f^ and Wnk1^d/d^ mice with implantation (green), without implantation (pink), and without implantation but presented embryos in the uterus (blue) on GD 4.5 and GD 5.5, n = 19 and 12 for Wnk1^f/f^ mice on GD 4.5 and GD 5.5, respectively; and n = 17 and 13 for Wnk1^d/d^ mice on GD 4.5 and GD 5.5, respectively. Red arrows indicate position of implantation sites. (E) Gross uterine morphology of Wnk1^f/f^ and Wnk1^d/d^ mice on GD 4.5, scale bars = 1 cm. (F) Hematoxylin and eosin staining of uterine cross sections at implantation site on GD 5.5 in Wnk1^f/f^ and Wnk1^d/d^ uteri, arrow indicates presence of maternal epithelium, and E = embryo. Scale bars = 100 µm. (G) Implantation marker *Lif* mRNA expression in the uteri as determined by qRT-PCR on GD 3.5 and GD 4.5 for Wnk1^f/f^ and Wnk1^d/d^ mice. Results shown are mean ± SD, * *p* < 0.05. (H) Expression of proliferative marker Ki-67 and implantation marker PGR on GD 4.5 in the stroma and epithelium of Wnk1^f/f^ and Wnk1^d/d^ mice. LE = luminal epithelium and S = stroma, scale bars = 25 µm.

We next examined whether the subfertile phenotype was associated with an implantation defect in the Wnk1^d/d^ mice. Dams were euthanized on GD 4.5, and embryo implantation was visualized by Evan’s blue dye staining. As expected, 84.2% of the control mice successfully permitted embryo implantation on GD 4.5 while the remaining 15.8% had no embryos present in the uterus as indicated by uterine flushing (Fig. 3.D, top panel and Fig. 3.E). On the other hand, significantly fewer (29.4%; *p* = 0.0019) of the mated Wnk1^d/d^ mice were able to form implantation sites on GD 4.5; however, 52.9% of the mice harboured fertilized embryos inside the uterus as identified by uterine flushing (Fig. 3.D, top panel and Fig. 3.E). Examination of the uterus on GD 5.5 showed 76.9% of the Wnk1^d/d^ mice with implantation sites which is, at this time point, comparable to their Wnk1^f/f^ control littermates (Fig. 3.D, bottom panel). Histological examination showed that the control Wnk1^f/f^ mice had already degraded the epithelium, enabling the embryo to invade the underlying stroma (Fig. 3.F, left), while Wnk1^d/d^ uteri had intact maternal epithelium at this time point with the embryo trapped inside the luminal space (Fig. 3.F). These findings demonstrate that in the Wnk1^d/d^ mice, implantation can still occur, but is severely delayed in comparison to Wnk^f/f^ mice.

In addition to the uterine expression, PGR is also expressed in the ovaries and pituitary, hence WNK1 was also ablated from those tissues. We next questioned whether the implantation phenotype could be attributed to a dysfunction of the ovary. Ovarian function was evaluated by assaying ovulation and ovarian steroid hormone levels. The mice were subjected to superovulatory regimen of gonadotropins, followed by mating to wildtype male mice, and euthanized on GD 1.5 to simultaneously monitor fertilization of the oocytes. We found a mild decrease albeit non-significant decrease in the number of 2-cell embryos produced by the Wnk1^d/d^ dams indicating that ovulation and fertilization was not affected (Fig. S2.A). Additionally, serum estradiol (E_2_) and progesterone (P_4_) levels were similar between the Wnk1^f/f^ and WNK1^d/d^ mice on GD 4.5 (Fig. S2.B and C), demonstrating that the ovaries were able to produce and maintain hormone levels. These results indicate that the main contributing factor for the delayed implantation was not a malfunction of the ovary.

Prerequisites for uterine receptivity are the production of leukemia inhibitory factor (LIF) from the uterine glands and the cessation of epithelial proliferation prior to implantation on GD 3.5 (23, 24); as well as suppression of epithelial PGR expression during implantation on GD 4.5 (25). Hence, we examined the uterus to see whether the delayed implantation was associated with impairment of those parameters. *Lif* gene expression on GD 3.5 was similarly induced in the Wnk1^f/f^ and Wnk1^d/d^ mice, which decreased to basal level on GD 4.5 in the Wnk1^f/f^ mice (Fig. 3.G). The Wnk1^d/d^ mice showed higher *Lif* levels on GD 4.5 (Fig. 3.G), however, this is unlikely to impact implantation. In the control mice, there was little to no expression of Ki-67 and PGR in the luminal epithelium on GD 4.5, as expected (Fig. 3.H). The Wnk1^d/d^ mice, however, maintained the expression of both proteins during the window of implantation (Fig. 3.G), indicating that there was a failure to impede epithelial proliferation. These findings illustrated that crucial implantation-associated molecular events were deregulated in the Wnk1^d/d^ mice.

### Abnormal embryo development and increased resorption in Wnk1^d/d^ mice

Interestingly, of the 29.4% mated Wnk1^d/d^ mice that were able to permit embryo implantation on time (GD 4.5), the number of implantation sites were similar to their Wnk1^f/f^ control littermates (Fig. 4.A). However, the number of implantation sites present on GD 5.5 was significantly lower in the Wnk1^d/d^ mice (Fig. 4.B and C). This finding indicated that the delay in implantation is associated with reduced number of implantation sites. Additionally, spacing between the implantation sites in the Wnk1^d/d^ mice were irregular whereas the implantation sites observed in the Wnk1^f/f^ mice were evenly distributed (Fig. 4.C). This is supported by a significant increase in the standard deviation of inter-implantation sites distance in Wnk1^d/d^ uteri compared to Wnk1^f/f^ uteri (Fig. 4.D). Interestingly, for the Wnk1^d/d^ mice that were able to implant promptly, implantation spacing was more evenly distributed (Fig. 3.E), suggesting that the delay may impact both implantation numbers and spacing. Examination of the uterus and embryo during mid-pregnancy (GD 8.5) further demonstrated that the Wnk1^d/d^ mice carried either resorbed embryos (Fig. 4.E, middle panel) or abnormally formed decidual balls (Fig. 4.E, right panel), compared to the normally sized decidual balls observed in the control mice (Fig. 4.E, left panel). Moreover, we also observed multiple embryos within one decidual zone (Fig. 4.E, right panel), possibly from the cluttered/delayed implantation. Morphology was evaluated by examining cross sections through the center of the decidual ball, which showed that the Wnk1^f/f^ mice have vascularized and initiated placentation (Fig. 4.F, black arrows and dashed lines, respectively), both of which were lacking in the Wnk1^d/d^ uteri. While this could be associated to delay in embryo development, it should not be ruled out that this may be a phenotype associated with the endothelial inhibition of WNK1 expression, as WNK1 is also known to function in endothelial cells (14, 26). Ultrasound scans demonstrated decreased gestation sac size (Fig. 4.G and H) and decreased embryo size at both GD 8.5 and GD 10.5 (Fig. 4.G and I). By GD 12.5, embryo resorption was frequently observed in the Wnk1^d/d^ mice (Fig. 4.G, bottom panel). Collectively, these findings demonstrate that uterine WNK1 ablation led to abnormal implantation and negatively impacted embryo development, resulting in the compromised pregnancy outcome and subfertility.

**Figure 4.**
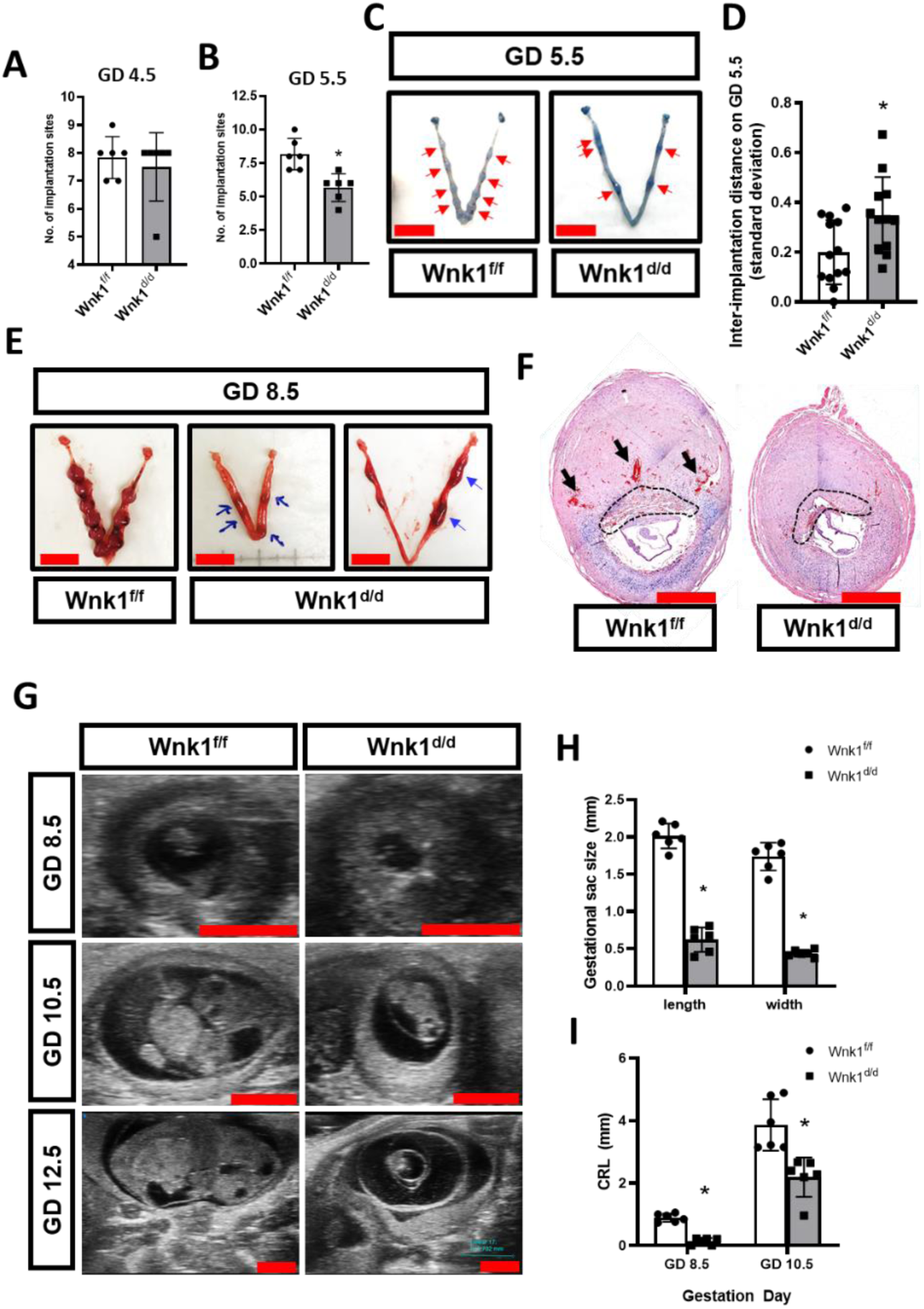
Abnormal embryo development and increased resorption n Wnk1^d/d^ mice. (A – B) Number of implantation sites on GD 4.5 (A) and GD 5.5 (B) in Wnk1^f/f^ and Wnk1^d/d^ mice (n = 6). (C, E) Uterine gross morphology on GD 5.5 (C) and GD 8.5 (E), with implantation sites on GD 5.5 marked by red arrows, and blue arrows indicate resorption and abnormal decidualization on GD 8.5. Scale bars = 1 cm. (D) Comparison of the standard deviation of inter-implantation distance in Wnk1^f/f^ and Wnk1^d/d^ mice (n= ≥ 12 uterine horns, 7 mice per genotype). (F) Hematoxylin and eosin staining of cross section through the centre of decidual ball on GD 8.5 from Wnk1^f/f^ and Wnk1^d/d^ mice, with black arrows and dashed line indicating vascularization and placentation, respectively. Scale bars = 2 mm. (G) Ultrasound scans of uterus and embryo during mid-pregnancy at GD 8.5, 10.5 and 12.5. Scale bars = 2 mm. (H and I) Quantification of gestational sac size by length and width on GD 8.5 (H), and embryo size by crown-rump length (CRL) on GD 8.5 and 10.5, as measured from ultrasound scans (I, n = 6). Results shown are mean ± SD, * *p* < 0.05.

### Loss of uterine WNK1 elevated AKT signaling

To fully characterize the molecular mechanisms underlying the loss of WNK1 induced-implantation defect, we next examined global gene expression profile by RNA sequencing (RNA-seq) in the uterus during receptivity. To ensure that the analysis was conducted only on the maternal uterine tissues and not the embryos, we used vasectomized wild-type males to induce pseudopregnancy in the Wnk1^f/f^ and Wnk1^d/d^ mice, which was confirmed by serum progesterone levels on pseudopregnancy day (PPD) 4.5 (Table S1). In total, there were 14,423 and 14,337 genes expressed in the Wnk1^f/f^ and Wnk1^d/d^ uterus, respectively; of which 14,024 were expressed in both. The transcriptomes were subjected to principle component analysis (PCA) as a measure of quality control, which segregated according to genotype indicating that the samples were well-characterized by genotype (Fig. S3). Using a defining threshold of *q*-value under 0.05 for significance and fold change (FC) over 1.5 as differential expression, we identified 1,727 significantly and differentially expressed genes (DEGs) in the Wnk1^d/d^ uterus during receptivity (Table S2). We then conducted detailed analyses to characterize the molecular alterations associated with uterine *Wnk1* ablation using the Database for Annotation, Visualization and Integrated Discovery (DAVID) bioinformatic database and Ingenuity Pathway Analysis (IPA). The top biological processes associated with the DEGs were adhesion, cell movement and locomotion, inflammation and blood vessel development (Table S3). Many important molecular functions associated with implantation were also deregulated in the Wnk1^d/d^ uteri, such as cell proliferation and apoptosis, Notch signaling, cell differentiation, epithelial to mesenchymal transition (EMT), cytokine production, and response to estrogen. Prediction of upstream regulator activity further showed altered activity for many important receptivity mediators, including the suppression of JAG, HEY2, PTEN and SERPINE1 (Fig. 5.A). On the other hand, TGFB1, ERBB2, AKT, estrogen, ERK, MUC1 and KLF5 were predicted to show increased activity (Fig. 5.A, for the complete list see Table S4). As WNK1 is a kinase, we next examined the alterations in the kinase phosphorylation network to understand the impact of WNK1 ablation. To this end, we employed an image-based phosphokinase array to simultaneously evaluate the phosphorylation status of multiple kinases in the uterus during receptivity (Fig. 5.B and C). Loss of WNK1 altered the phosphorylation of various kinases, including TOR (mTOR), SRC, PRAS40 (AKT1S1), JNK, AMPKα1 (PRKAA1), GSK-3α/β (GSK3A and GSK3B) and AKT (Fig. 5.B and C, all kinases with > 2 fold change in phosphorylation are shown in Fig. S4.A). The phosphorylation of AKT, GSK-3α/β and PRAS40 were independently validated via western blotting (Fig. S4.B) and all showed elevated phosphorylation in Wnk1^d/d^ uteri during receptivity. Interestingly, AKT was identified as an activated upstream regulator by IPA, and the phosphokinase array demonstrated its elevated phosphorylation in the Wnk1^d/d^ uteri during receptivity (Fig. 5.B, C and Fig. S4.B). Furthermore, elevated AKT phosphorylation on GD 4.5 was confirmed in both the epithelium and the stroma (Fig. 5.D). Indeed, we found that in the control mice, phosphorylation of AKT was actively suppressed as the mice transitioned into the receptive phase from GD 3.5 to PPD 4.5, however, the Wnk1^d/d^ mice maintained high AKT phosphorylation both prior to and during receptivity (Fig. 5.E).

**Figure 5.**
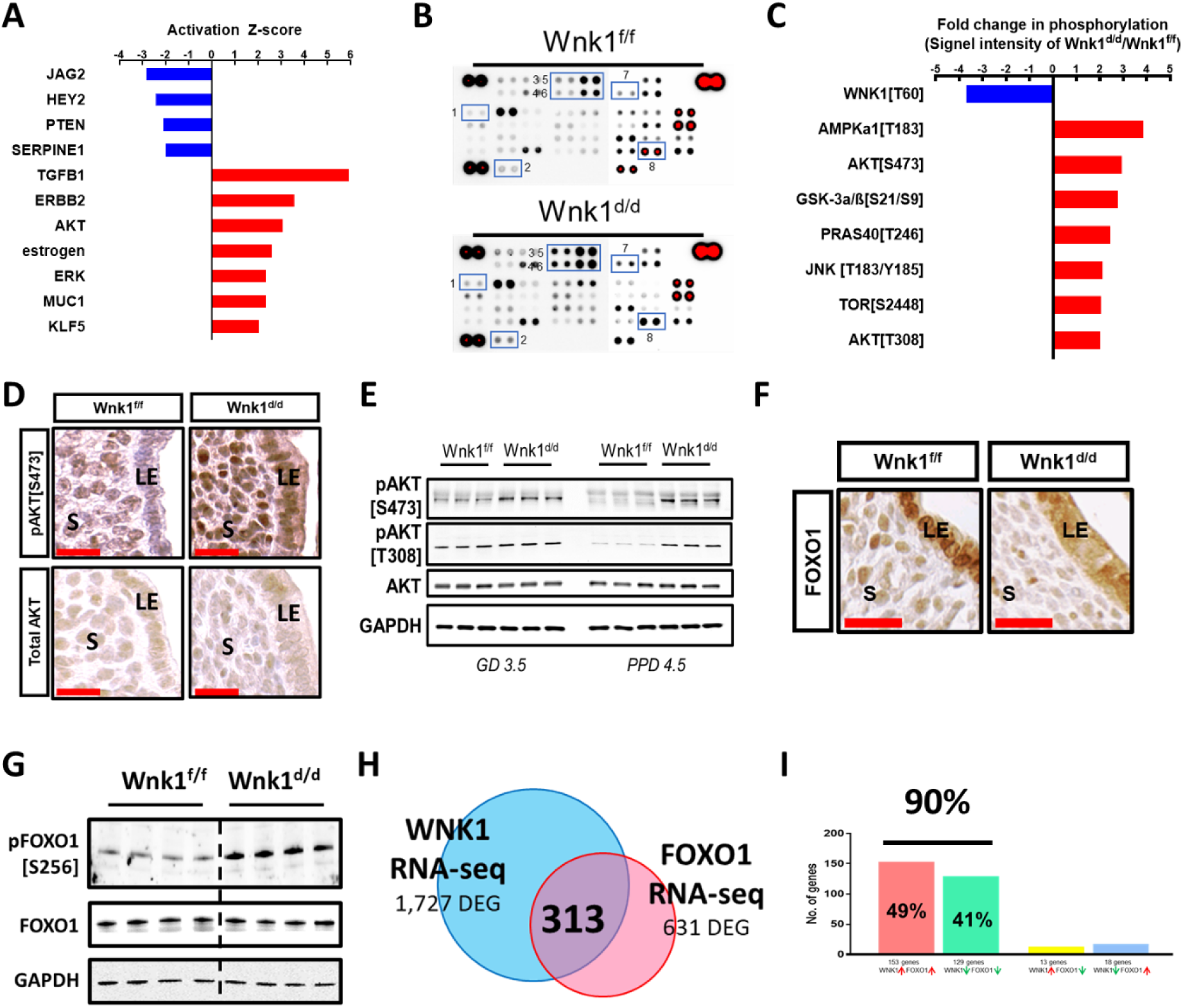
Loss of uterine WNK1 elevated AKT signaling. (A) Activity of upstream regulators as predicted by Ingenuity Pathway Analysis (IPA) based on the altered uterine transcriptome of Wnk1^d/d^ mice on PPD 4.5. See Table S4 for complete list. (B and C) Kinome phosphorylation status in Wnk1^d/d^ and Wnk1^f/f^ uteri on pseudopregnancy day (PPD) 4.5, with selected alterations shown in (C). All kinases with > 1.5 fold change in signal intensity as quantified by ImageJ is shown in Fig. S4. Results were acquired using pooled uterine lysate from 6 mice in each group. (D and E) Expression of phosphorylated and total AKT in Wnk1^f/f^ and Wnk1^d/d^ uteri on GD 4.5 as shown by immunohistochemistry ((D), LE = luminal epithelium and S = stroma), and on GD 3.5 and PPD 4.5 as shown by Western blotting (E), scale bars = 25 µm. (F) Expression of AKT-regulated implantation marker FOXO1 on GD 4.5 in the stroma and epithelium of Wnk1^f/f^ and Wnk1^d/d^ mice. LE = luminal epithelium and S = stroma, scale bar = 25 µm. (G) Western blot analysis showing levels of phosphorylated and total FOXO1 in Wnk1^f/f^ and Wnk1^d/d^ uteri on PPD 4.5. (H) Comparison of differentially expressed genes (DEGs) between the uteri of WNK1 ablated mice vs. their control littermates (1727 DEG; blue) and FOXO1 ablated mice vs. their control littermates (611 DEG; pink) identified 313 common genes. (I) Percentage of the 313 genes categorized into commonly upregulated (pink), commonly downregulated (green), or upregulated in one and downregulated in the other (yellow and blue).

We have shown earlier that implantation is compromised in the Wnk1^d/d^ mice, and the AKT-regulated transcription factor FOXO1 is an indispensable mediator of implantation, as mice lacking uterine FOXO1 expression suffer infertility due to failed implantation (27). Interestingly, AKT is known to exert an inhibitory effect on FOXO1 transcriptional activity via directly phosphorylating it, resulting in its nuclear exclusion (28, 29). To examine whether loss of WNK1 is impairing implantation through AKT and FOXO1, we examined FOXO1 expression using immunohistochemistry, and found that indeed, there was a marked decrease in its nuclear form in both the luminal epithelium and underlying stroma (Fig. 4.F). This is further confirmed by the increase in the levels of phosphorylated FOXO1 in the Wnk1^d/d^ uteri (Fig. 4.G). As FOXO1 nuclear exclusion prevents its transcriptional activity, we compared WNK1-regulated genes and FOXO1-regulated genes in the uterus during the receptive phase (27). This revealed that roughly half of the FOXO1-regulated genes are also deregulated in Wnk1^d/d^ uteri (Fig. 5.H), and strikingly, 90% of those genes were deregulated in the same direction under WNK1 and FOXO1 deficient conditions – these included known implantation and decidualization associated genes such as *Msx2, Wnt5a* and *Muc1* (Fig. 5.I, table S2, and Vasequez *et al*., 2018 (27, 30-32)). These findings indicate that uterine loss of WNK1 led to elevated AKT phosphorylation and signaling, which was evident through the increased FOXO1 phosphorylation and nuclear exclusion. This in turn, led to deregulation of FOXO1-regulated genes during receptivity. Additionally, it appears that the impact of WNK1 on both AKT and FOXO1 is maintained in the same way in both the epithelial and stromal compartments, as both responded similarly to WNK1 deficiency.

### WNK1 regulates FOXO1 localization via AKT, which is associated with decreased PP2A expression and activity

Having demonstrated that the loss of WNK1 led to increased phosphorylation of AKT and FOXO1 in mouse uteri, we next examined whether this regulatory axis was similarly maintained in human endometrial HEC1A (epithelial) and THESC (stromal) cells. Using small interfering RNA against *WNK1* (siWNK1), WNK1 protein expression was inhibited, which robustly induced AKT and FOXO1 phosphorylation in both cell lines (Fig. 6.A). In order to test whether AKT facilitated FOXO1 localization downstream of WNK1, we next treated these cells with an AKT inhibitor, GDC0941, and examined whether it could rescue WNK1 ablation-induced phosphorylation and nuclear exclusion of FOXO1. FOXO1 localization clearly decreased in the nucleus of both cells after transfection with siWNK1 (Fig. 6.B, panels 1 VS 2, and 4 VS 5). However, when the siWNK1 transfected cells were treated with GDC0941, nuclear FOXO1 was readily restored (Fig. 6.B, panels 3 and 6). This suggested that WNK1 inhibition-induced nuclear exclusion of FOXO1 is mediated through AKT. This is further supported by the findings that AKT inhibition rescued WNK1 knock-down induced FOXO1 phosphorylation (Fig. 6.C). Interestingly, GDC0941 treatment reduced the phosphorylation of AKT and FOXO1 to a level that is lower than seen in the siCTRL transfected, untreated cells (considered basal level). As GDC0941 inhibits AKT through its upstream regulator PI3K (33), it is likely that PI3K lies upstream of WNK1 in regulating AKT. Indeed, none of the PI3K family members were impacted by WNK1 inhibition, including p110-α, p110-β, p110-γ, Tyr458 phosphorylated p85 and Tyr199 phosphorylated p55 (Fig. 6.C). Similarly in mice, the expression of these proteins in the uterus were comparable between the Wnk1^f/f^ and Wnk1^d/d^ mice during receptivity (Fig. 6.D).

**Figure 6.**
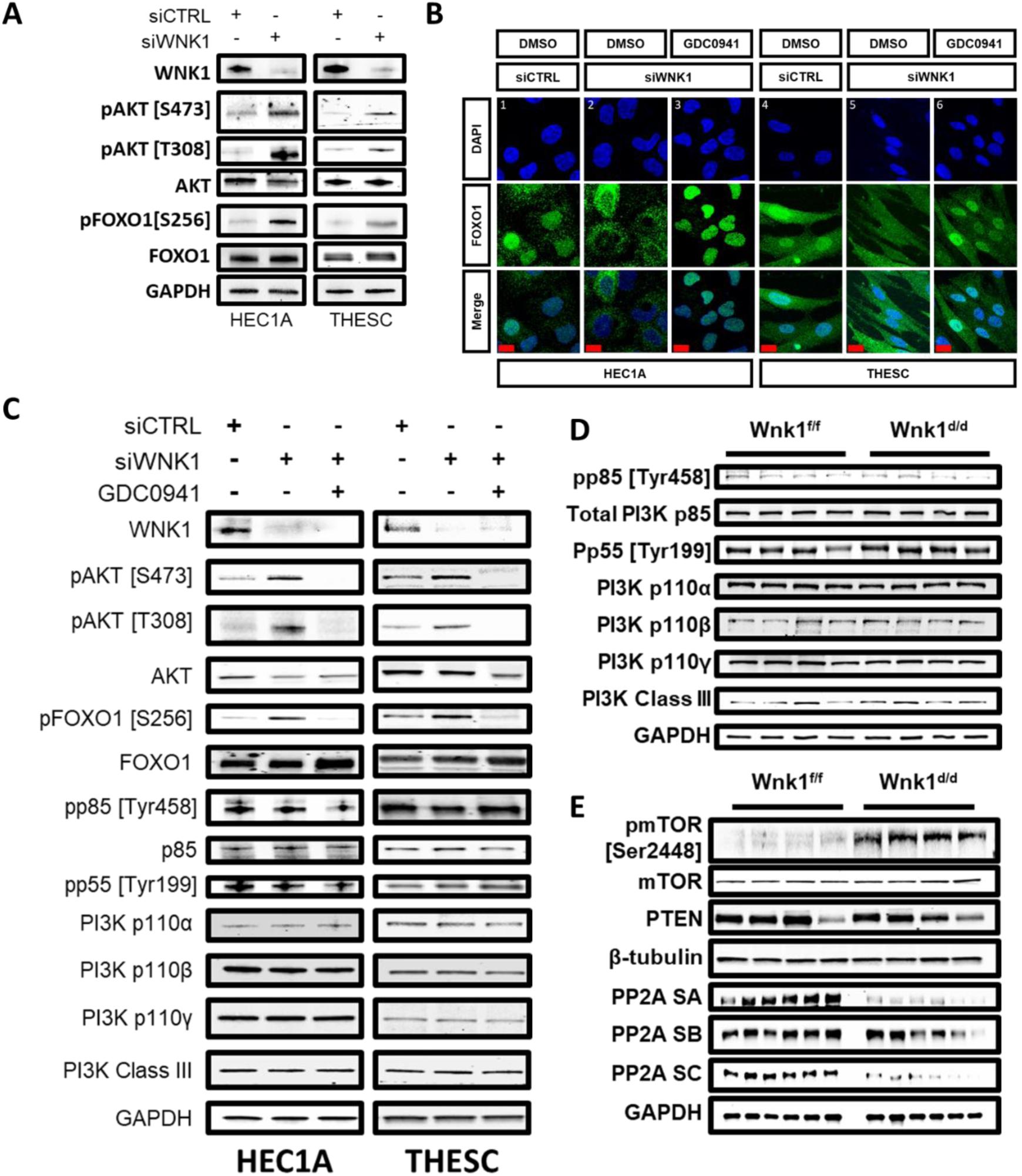
WNK1 ablation led to FOXO1 nuclear exclusion via AKT phosphorylation which is associated with decreased PP2A phosphatase expression. (A) Western blot showing levels of phosphorylated and total AKT and FOXO1 in HEC1A and THESC cells transfected with 24 nM siCTRL or siWNK1. (B) Immunofluorescence showing FOXO1 subcellular localization (green), with nuclei presented in DAPI in HEC1A and THESC control cells (1, 4), siWNK1 transfected cells (2, 5), and GDC0941 treated, siWNK1 transfected cells (3, 6), scale bars = 20 µm. (C) Expression of FOXO1, AKT and PI3K members in HEC1A and THESC cells transfected with siCTRL, siWNK1, and treated with AKT inhibitor GDC0941. (D and E) Expression of PI3K proteins (D), and mTOR, PP2A subunits and PTEN (E) in Wnk1^f/f^ and Wnk1^d/d^ uteri on PPD 4.5.

We then explored the possible mechanisms through which WNK1 could regulate AKT phosphorylation and activity. A search of the upstream regulators predicted by IPA identified several candidates with altered activities in the Wnk1^d/d^ uteri, including PTEN, PPP2CA, and sirolimus (rapamycin, Table S4). PTEN and PPP2CA are both phosphatases that regulate AKT phosphorylation, and both displayed repressed activities in the Wnk1^d/d^ mice during receptivity (Z-scores of -2.079 and -1.195, respectively, Table S4). Sirolimus, on the other hand, is a drug targeting the kinase mTOR, which was strongly inhibited (Z-score of -2.95, Table S4). We found that mTOR phosphorylation and PP2A subunits A and C were altered in the Wnk1^d/d^ mice, while PTEN level was not significantly different (Fig. 6.E). This finding suggested that increased AKT phosphorylation in the Wnk1^d/d^ mice may be mediated through elevated mTOR or repressed PP2A activity. As mTOR is both a regulator and a substrate of AKT (34, 35), we examined whether WNK1 ablation-induced AKT phosphorylation is mediated through mTOR. We inhibited mTOR activity using rapamycin and examined AKT/FOXO1 phosphorylation as well as FOXO1 localization as a readout of AKT activity. As shown in Fig. S5.A, rapamycin treatment did not reverse the nuclear exclusion of FOXO1 induced by WNK1 inhibition. Additionally, AKT and FOXO1 phosphorylation was not rescued by rapamycin treatment (Fig. S5.B). Similar results were observed in HEC1A cells where WNK1 and mTOR double knock-down failed to rescue AKT and FOXO1 phosphorylation (Fig. S5.C). Thus, mTOR is likely not the WNK1 mediator controlling AKT activity, and its elevated phosphorylation is a result of elevated AKT activity, rather than its cause.

### WNK1 regulates AKT phosphorylation through direct interaction with PPP2R1A

So far, we have shown that WNK1 ablation led to subfertility in mice which is associated with an array of abnormalities including impaired implantation, altered morphology, epithelial hyperplasia and adenomyosis. Upstream regulator prediction and validation experiments identified elevated AKT phosphorylation in the Wnk1^d/d^ uteri. This elevation was demonstrated to have functional consequences, evident through the deregulated FOXO1 transcriptional activity. In addition, there was a concomitant repression of the negative AKT regulator, PP2A. We explored the possible regulatory link between WNK1 and PP2A/AKT using a non-biased WNK1 immunoprecipitation-mass spectrometry (IP-MS) approach to identify WNK1 binding partners. Successful WNK1 IP was confirmed by examining the lysate for WNK1 expression after immunoprecipition using a rabbit IgG (negative control) or WNK1 targeting antibody from HEC1A cells (Fig. S6), and the peptides identified by mass-spectrometry are listed in Table S5. Amongst those were peptides belonging to WNK1 itself, as well as a known WNK1 substrate, oxidative stress responsive kinase 1 (OXSR1/OSR1) (12), confirming the validity of the pull-down results (Table S5).

Putative WNK1 binding proteins identified in this experiment included Wnt regulators (OFD1 and CCDC88C), chromosome modulating and DNA repair proteins (SMCA1, KIF11, FANCI, RAD50 and SLC25A5), proteins associated with the endoplasmic reticulum and ribosomal functions (UGGT1, SEC23A, HYOU1, EMC1, AIFM1, HM13, SCFD1) as well as the mitochondria (AIFM1, SLC25A5). Of particular interest were the components of protein phosphatase complexes PP2A (PPP2R1A) and PP6 (PPP6R3), as both regulate AKT signaling (36, 37). Since the enzymatic activity of PP2A was predicted by IPA as repressed in the Wnk1^d/d^ mice during receptivity (PPP2CA, Table S4), we postulated that these observations were associated with the interaction between WNK1 and PPP2R1A, the alpha isoform of the scaffold subunit A of PP2A. In order to confirm the interaction of WNK1 and PPP2R1A, a YFP-tagged WNK1 (c4161, Fig. S7) was expressed in HEC1A cells, then immunoprecipitated using a YFP nanobody, followed by detection for PPP2R1A in the pulldown. We first confirmed that c4161 transfection induced exogenous WNK1 expression when compared to the control cells transfected with YFP only expressing construct (cYFP, Fig. 7.A). WNK1 was subsequently detected in the lysate immunoprecipitated for YFP (Fig. 7.B, upper panel), which co-immunoprecipitated with PPP2R1A (Fig. 7.B, middle panel).

**Figure 7.**
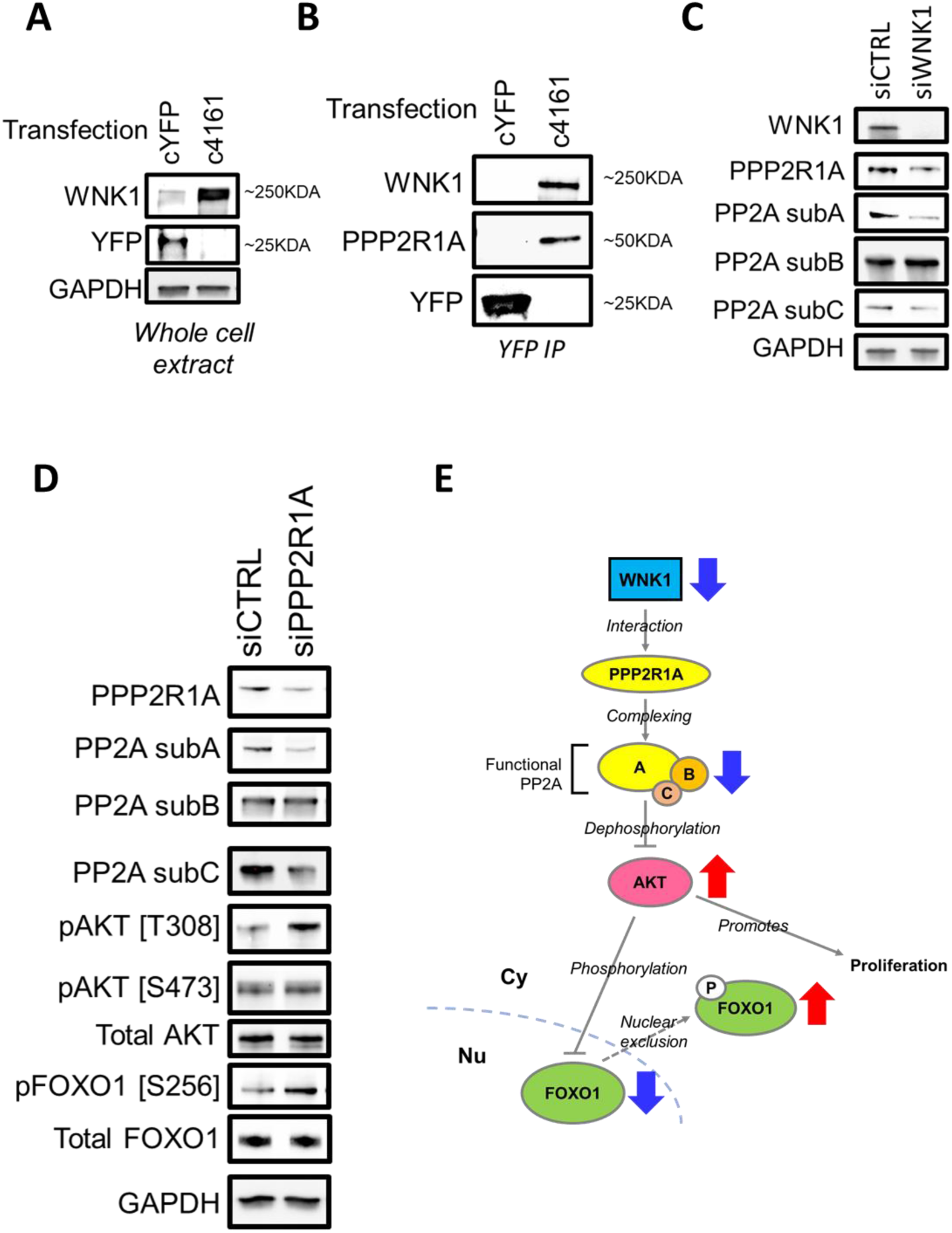
WNK1 regulates AKT signaling through direct interaction with PPP2R1A. (A) WNK1 and YFP expression in HEC1A cells transfected the YFP expressing control plasmid (cYFP) or YFP-tagged WNK1 expression construct (c4161). (B) Co-immuniprecipitation of WNK1 and PPP2R1A with YFP in HEC1A whole cell lysate, as indicated by western blotting. (C) Expression of PPP2R1A and PP2A subunits in HEC1A cells transfected with 24 nM siCTRL or siWNK1 for 72 hours. (D) Expression of PP2A subunits A, B and C, AKT and FOXO1 in HEC1A cells transfected with 72 nM siCTRL or siPPP2R1A for 72 hours. (E) Diagram illustrating the WNK1-PP2A-AKT-FOXO1 signaling axis. WNK1 physically interact with PPP2R1A, the alpha isoform of the scaffold subunit that forms the functional PP2A subunit. PP2A negatively regulates AKT, and AKT negatively regulates FOXO1 by phosphorylation and nuclear exclusion. AKT also promotes epithelial cell proliferation. As indicated by the blue and red arrows, decreased or loss of WNK1 will then lead to decreased PP2A activity, AKT hypersignaling and increased cytoplasmic FOXO1 retention and epithelial proliferation.

Having confirmed the WNK1-PPP2R1A interaction, we next explored the biological implications of this interaction. The PP2A phosphatase complex is comprised of the scaffold subunit A with 2 isoforms, the regulatory subunit B with 13 isoforms and the enzymatic subunit C with 2 isoforms. As shown earlier, uterine WNK1 ablation led to decreased protein levels of subunits A and C (Fig. 6.E), yet RNA-seq showed no alteration in transcription of the 4 genes encoding these 2 subunits (*Ppp2ca, Ppp2cb, Ppp2r1a* and *Ppp2r1b*) in the Wnk1^d/d^ mice. It has been reported that the stability of the PP2A subunits depends on their association with each other (38). Hence, reduced subunit levels could be an indication that the complexing capacity of the subunits were reduced in the absence of WNK1, leading to their degradation. We therefore postulated that the WNK1-PPP2R1A interaction is necessary for the association of the PP2A subunits. To test this idea, we examined the levels of PPP2R1A, total PP2A subunit A and total PP2A subunit C in WNK1 knock-down HEC1A cells, and accordingly found their reduced levels upon WNK1 inhibition (Fig. 7.C). Lastly, to validate that PP2A mediates AKT/FOXO1 signaling, we inhibited PPP2R1A expression in HEC1A cells using siRNA, and examined the components of the PP2A-AKT-FOXO1 signaling axis. As expected, PPP2R1A knock-down caused a reduction in total subunits A and C of PP2A (Fig. 7.D). Interestingly, AKT phosphorylation was selectively induced on threonine 308, but not serine 473 after PPP2R1A knock-down (Fig. 7.D). This nonetheless, translated to elevated FOXO1 phosphorylation, indicating that loss of PP2A activity-induced AKT phosphorylation on this residue alone is sufficient to increase FOXO1 phosphorylation (Fig. 7.D). These findings confirmed that in endometrial cells, WNK1 controls AKT activity through modulating its phosphorylation, which is partially mediated through PP2A (Fig. 7.E). As such, loss of WNK1 led to decreased PP2A activity and increased AKT phosphorylation, resulting in the pathological outcomes associated with AKT hypersignaling such as hyperplasia and FOXO1 deregulation (Fig. 7.E, blue and red arrows).

## Discussion

Reproductive biology has relied profoundly on transcriptomic analyses to identify novel players that may serve crucial functions in the regulation of fertility. While this approach has uncovered many key components in the reproductive tract, it is unable to detect alterations at the proteomic level, such as post translational modifications (PTMs). In many cases, the PTMs control protein activity and stability, and hence are the actual determinants of functional output. Using a proteomic approach, we identified WNK1 as a potential regulator of uterine biology with previously unreported reproductive functions.

In this study, we examined the *in vivo* function of WNK1 using a whole-uterus WNK1 knock-out mouse model. We demonstrate that loss of WNK1 led to hyperplasia, adenomyosis and impaired implantation, which could all negatively impact fertility. Importantly, we demonstrate for the first time that normally, WNK1 robustly represses AKT activity and that loss of WNK1 led to increased AKT phosphorylation and signaling. This was evident through the increased mTOR and FOXO1 phosphorylation (28, 29, 35), resulting in nuclear exclusion of the latter and disrupted embryo implantation (27). It is worth noting that the Wnk1^d/d^ mice only partially recapitulated the uterine FOXO1 knock-out phenotype (27), likely due to reduced FOXO1 activity rather than a complete inhibition. Additionally, AKT is well known for promoting cell proliferation and has long been explored as a target for therapeutic purposes in anti-cancer treatment (39). This is consistent with our observation that Wnk1^d/d^ mice displayed epithelial hyperplasia resulting from escalated proliferation. Evidence has also shown a link between adenomyosis and estrogen induced AKT overactivity (40). Although not the focus of this work, the cellular changes and molecular events associated with WNK1 deficiency induced adenomyosis seems to parallel observations made in humans, including excessive ECM, elevated *Moesin* expression and AKT hypersignaling (20, 22). This indicates that the Wnk1^d/d^ mice could serve as an ideal model system to study adenomyosis in the future. Interestingly, there are evidence in literature showing that WNK1 is a substrate of AKT in other cellular and animal systems (41, 42). We did notice a reduction in phosphorylated WNK1 after AKT inhibition in both HEC1A and THESC cells (data not shown), suggesting that AKT could reciprocally interfere with WNK1 activity. These, together with our demonstration that in endometrial cells, WNK1 inhibits AKT phosphorylation and activity suggest that the WNK1-AKT relationship involves a negative feedback loop and is likely more complex than previously thought.

Given that 30% of the Wnk1^d/d^ mice were able to implant promptly with normal numbers of embryos, we rarely observed normal sized litters from these mice. This suggested that even after a seemingly normal implantation, there must exist other impairments in subsequent pregnancy development accounting for the compromised fertility. This is supported by various observations during the breeding trial. The significant proportion of Wnk1^d/d^ mice succumbing to pregnancy complications, including death near term and dystocia which indicated poor support of pregnancy and impairment in uterine muscle contractility. This could be attributed to impaired decidualization – indeed, our previous *in vitro* study demonstrated that WNK1 is a regulator of decidualization (3). We did not extensively characterize decidualization in this study due to the preceding implantation defect – which complicates decidualization data interpretation. However, transcriptomic analysis did identify alterations in several decidualizing regulators including Notch (HEY2 and JAG2), ERK and MUC signaling (43-45). Therefore, we are quite confident in speculating that loss of WNK1 will likely negatively impact decidualization *in vivo*. Additionally, the premature loss of fertility in those mice that survived to the end of the breeding trial suggested that the postpartum tissue repair and remodeling may also be impacted by loss of WNK1. Interestingly, Zhu *et al*. reported on the AKT dependent endometrial stromal cell repair in humans (46), providing a possible explanation for the premature sterility.

We further characterized the regulatory mechanism linking AKT to WNK1 and identified PP2A as an intermediate. We found that loss of WNK1 reduced PP2A subunits A and C in both humans and mice endometrial cells, and reduced PP2A phosphatase activity. Interestingly, RNA-seq results did not showed that the 4 genes encoding subunits A and C were altered. This suggests that the altered protein levels may be a result of altered protein stability rather than expression. It has been previously reported that the half-life of the PP2A subunits is dependent on their ability to complex with each other (38). Therefore, a possibility is that WNK1 facilitates the binding of the subunits, resulting in PP2A complex stabilization. As such, in the absence of WNK1, the decreased coupling of the subunits to each other led to shorter half-lives and decreased protein levels. It is worth noting that the antibody against PP2A subunit B used in this study targets only one of the thirteen isoforms (PPP2R2A). While this particular isoform was unaffected by loss of WNK1, it is possible that other subunit B isoforms may be affected. Mechanistically, WNK1 directly interacts with PPP2R1A and hence this interaction may be crucial for PP2A complex formation, however, further experimentation will be necessary to test this hypothesis. Future studies will aim to understand the biochemical nature of the PPP2R1A-WNK1 interaction, for example, whether WNK1 phosphorylates PPP2R1A, and if so, whether the phosphorylation interferes with PP2A subunits coupling. It is worth noting that there must be another mechanism through which WNK1 is repressing AKT, as PPP2R1A knock-down in human cells restored only the phosphorylation on threonine 308. However, WNK1 ablation induced phosphorylation on both threonine 308 and serine 473, indicating that other regulators must be involved.

As aforementioned, uterine WNK1 ablation exhibited pleiotropic effects including epithelial hyperplasia and adenomyosis. Although neither is cancerous, both are progressive conditions which may lead to malignant transformation (47, 48). Functional interpretation of the transcriptome reiterated this, where many cancer development and progression associated signaling pathways were altered in the Wnk1^d/d^ uteri, including elevated TGFB, AKT and estrogen (48-51). Strikingly, a recurrent mutation of *Ppp2r1a* is associated with serous endometrial carcinoma (52, 53), and this mutation has been found to impact oncogenic signaling through a dominant negative effect (54). It is possible that WNK1 could act to protect cells against cancer progression through stabilizing PP2A subunits and hence activity. While this speculation is made based on data from our current study, there are also studies suggesting that WNK1 promotes oncogenesis through stimulating angiogenesis (55). WNK1’s exact role in endometrial cancer is yet unknown but worth further exploring, as mutation of the *WNK1* gene is an existing condition in humans. Expanding our knowledge on the potential pathogenic or beneficial consequences associated with altered WNK1 expression and regulation could be a valuable insight to have in the development of therapeutic interventions in the future. As the existing condition in humans is a gain-of-function mutation, one future direction is to characterize the effect of WNK1 over-expression on AKT signaling in the female reproductive tract, and the implication on uterine biology and oncogenesis. In summary, this study is a first to explore the reproductive function of WNK1 *in vivo*. We demonstrate that WNK1 is critical in maintaining normal uterine morphology, mediating epitheial homeostasis and implantation.

## Methods

### Study approval

This study was conducted according to the federal regulations regarding the use of human subjects. Procedures were approved by the following ethics committee: Institutional Review Board/Committee-A (IRB) of Greenville Health System under IRB file #Pro0000093 and Pro00013885 and the University of Chapel Hill at North Carolina IRB under file #: 05-1757. Written, informed consents were obtained from all patients prior to participation.

All animal studies were conducted in accordance with the Guide for the Care and Use of Laboratory Animals, as published by the National Institute of Health. Animal protocols were approved by the Animal Care and Use Committee (ACUC) of National Institute of Environmental Health Sciences (protocol numbers 2015-0012 and 2015-0023). The mice were housed with a maximum of 5 per cage with a 12-hour light and dark cycle, and fed irradiated Teklad global soy protein-free extruded rodent diet (Harlem Laboratories, Inc., Indianapolis, IN) and fresh water ad libitum. Euthanization was carried out by carbon dioxide inhalation followed by cervical dislocation.

### Generation of transgenic mice

The Wnk1^f/f^ mice were a kind gift from Dr CL Huang (University of Iowa Healthcare), the generation of which was described in a previous publication(14). Briefly, the *Wnk1* floxed allele was established by insertion of *loxP* sites into the 5’ and 3’ region of exon 2 of the mouse *Wnk1* gene. The Wnk1^f/f^ mice were crossed to mice carrying Cre under the control of the progesterone receptor (PGR^Cre^) to generate conditional uterine Wnk1 ablated mice (Wnk1^d/d^, Fig. S1) (16).

### Fertility trial

Seven-week old Wnk1^f/f^ (control) and Wnk1^d/d^ (experimental) mice were housed with wild-type CD1 males for a period of 6 months, during which the mice were monitored daily for pregnancy and delivery. Upon the first observation of delivery, the total number of pups including both live and dead were recorded.

### Implantation determination and pseudopregnancy

Mice of age 6-10 weeks were housed with wild-type C57BL/6J males, and monitored each morning until vaginal plug is observed (indicating that mating has occurred). The first noon following which the vaginal plug was seen was defined as gestation day (GD) 0.5. Mice were anesthetized by isoflurane inhalation on GD 4.5 and 5.5, followed by retro-orbital administration of 200 µL of 1% Evans Blue Dye prepared in phosphate buffered saline (PBS). Mice were then euthanized, and the uterine horns harvested for implantation determination and imaging. For mice sacrificed on GD 4.5, the uterine horns were flushed using a p200 pipette if no implantation sites were observed, and the eluant was examined under brightfield microscope to determine presence of blastocysts. The uterine horns were then fixed 48 hours in 4% paraformaldehyde (PFA) prepared in PBS for histology and immunohistochemstry, or frozen on dry ice for RNA and protein extraction. For RNA-seq, pseudopregnancy was induced by mating the females to vasectomized wild-type male mice, and all procedures were conducted as described above except for the Evans Blue Dye injection, as no implantations were expected.

### Superovulation assay

Three-week old Wnk1^f/f^ and Wnk1^d/d^ mice were subjected to a superovulation regimen, which began with intraperitoneal administration of 5 IU of pregnant mare’s serum gonadotropin (catalog no. 493-10-2.5, Lee Biosolutions), followed by 5 IU of human chorionic gonadotropin (catalog no. 869031, EMD Millipore) 48 hours later. Superovulated mice were placed with wild-type CD1 males overnight. Mating was confirmed by presence of vaginal plug the next morning (GD 0.5), and mice were euthanized on GD 1.5 followed by oviduct flushing. The number of embryos was determined by counting under a brightfield microscope.

### Serum collection

On GD or pseudopregnancy day (PPD) 4.5, mice were anesthetized by intraperitoneal administration of Fetal Plus (1mg/10g body mass) and whole blood was collected via retro-orbital puncture. Blood was allowed to clot at room temperature for approximately 30 minutes, then centrifuged at 1000 X G for 10 minutes at 4°C. The supernatant (serum) was moved into a fresh tube and stored at -80°C until hormone testing. Hormone testing was conducted by the Ligand Core Laboratory of University of Virginia, Center for Research in Reproduction.

### High frequency ultrasound imaging

On GDs 8.5, 10.5 and 12.5, high frequency ultrasound imaging was used to evaluate the uterus and embryo development. Dams were anesthetized by isoflurane inhalation and placed onto an electric heating pad to maintain body temperature. Abdominal hair was removed using depilatory cream (Nair_TM_ Church & Dwight Co. Trenton, NJ), and eye lubricant was applied to prevent desiccation. Dams were manipulated into a supine position for the scan while heart rate and body temperature were continuously monitored. Images were visualized and captured using the VisualSonics VevoR 2100 Imaging System with a 550s scan head (Fujifilm VisualSonics Inc., Toronto, ON) at 55 megahertz. Each scanning session was limited to maximum 15 minutes, after which the dams were monitored until full recovery.

### Tissue processing, histology, immunohistochemical and immunofluorescence staining

For histology, immunohistochemistry and immunofluorescence, the tissues were similarly processed as described below. After 48 hour fixation, tissues were placed into 70% ethanol for a minimum of 48 hours. Tissues were then dehydrated and embedded in paraffin blocks and sectioned to 5 µm thickness onto glass slides. Slides were heated at 60°C for 10 minutes, followed by 5 minutes cooling. Sections were deparaffinized by 3 serial incubations in Citrisolv clearing agent (catalog no. 22-143-975, Thermo Fisher), followed by rehydration through decreasing % of ethanol. For histology, sections were subjected to hematoxylin and eosin (H&E) and Masson’s trichrome staining, followed by dehydration through increasing % of ethanol, incubation in Citrisolv and coverslipping. For immunohistochemistry, sections were subjected to antigen retrieval after rehydration by boiling in the Vector Labs Antigen Unmasking Solution as per manufacturer’s instructions (H-3300, Vector Laboratories, Burlingame, CA, USA). Blocking of endogenous peroxidase was performed by treating the sections with 3% hydrogen peroxide diluted in distilled water for 10 minutes at room temperature. Tissues were blocked in 5% normal donkey serum (NDS) for 60 minutes at room temperature, prior to overnight incubation with the primary antibody at 4°C. The slides were washed twice in PBS for a total of 10 minutes at room temperature and secondary antibody diluted in 1% w/v bovine serum albumin (BSA) prepared in PBS was applied. The ABC reagent was applied to tissue according to the manufacturer’s instructions (Vector Labs ABC PK-6100, Vector Laboratories). Signals were developed using the Vector Labs DAB ImmPACT Staining Kit (Vector Labs SK-4105, Vector Laboratories). Finally, the tissues sections were counterstained with hematoxylin and dehydrated through increasing ethanol concentration, followed by Citrisolv incubation and coverslipping. For immunofluorescence, tissue sections were subjected to antigen retrieval as described above. Tissues were blocked in 0.4% v/v Triton X-100, 1% BSA and 5% NDS for 30 minutes at room temperature followed by overnight incubation in primary antibody prepared in 0.4% Triton X-100/PBS at 4°C. Sections were washed 3 times 5 minutes in PBS and incubated with secondary antibodies diluted in 0.4% Triton X-100/PBS for 90 minutes at room temperature. Finally, the slides were washed 3 times 5 minutes in PBS, and coverslipped in DAPI containing mounting medium (Vectorshield Hardset™ Antifade Mounting Medium, catalog no. H-1400, Vector Laboratories). Details of antibodies used in this study are provided in table S6.

### RNA extraction and cDNA conversion

The frozen tissues were disrupted in TRIzol reagent (Thermo Fisher) by bead milling, followed by 2 aqueous phase separations using 1-Bromo-3-chloropane and chloroform. Pure ethanol was added to the aqueous layer, and the RNA was extracted using the Qiagen RNEasy RNA mini prep kit columns as per manufacturer’s instructions (Qiagen, Valencia, CA). Resulting RNA concentration and quality was determined using the NanoDrop ND-1000. cDNA was generated by reverse transcription using the M-MLV Reverse Transcriptase (catalog number 28025013 Thermo Fisher) following the manufacturer’s instructions.

### qRT-PCR

qRT-PCR was performed using the SsoAdvancedTM Universal SYBR Green Supermix (1725274, Bio-Rad) with the following primers (from 5’ to 3’, F = forward and R = reverse): *Wnk1 –* AGGCAGAGATTCAAAGAAGAGG (F) and CCCAGGAATCATAGAATCGAACA (R); *Msn* – CCATGCCGAAGACGATCA (F) and CCAAACTTCCCTCAAACCAATAG (R); and *Foxa2* – GAGACTTTGGGAGAGCTTTGAG (F) and GATCACTGTGGCCCATCTATTT (R). *Lif* expression was determined using the Taqman Master Mix (Life Technologies) and Taqman probes (Applied Biosystems). The Delta delta Ct values were calculated using 18S RNA control amplification results to acquire the relative mRNA expression for each sample.

### RNA-sequencing

For each mouse, 1 µg of total uterine RNA was sent to the NIH Intramural Sequencing Center to create a libarary using the TruSeq RNA Kit (Illumina, San Diego, CA, USA) following the manufacturer’s instructions. The RNA libraries were sequenced with a HiSeq 2000 System (Illumina). The raw RNA reads (75 nt, paired-end) were processed by filtering with average quality score greater than 20. Reads that passed the initial processing were aligned to the mouse reference genome (mm10; Genome Reference Consortium Mouse Build 38 from December 2011) using TopHat version 2.0.4 (56). Expression values of RNA-seq were expressed as fragments per kilobase of exon per million fragments (FPKM). Differential expression was calculated using Cuffdiff function from Cufflinks version 2.2(57). Transcripts with the average FPKM > 1 in at least one group, *q*-value < 0.05 and at least 1.5-fold difference in FPKM were defined as differentially expressed genes (DEGs). The data discussed in this publication have been deposited in NCBI’s Gene Expression Omnibus and are accessible through GEO Series accession number GSE144802. Functional annotation for the differentially expressed genes derived from RNA-seq were analyzed by Ingenuity Pathway Analysis (IPA) and Database for Annotation, Visualization, and Integrated Discovery (DAVID)(58).

### Human phospho-kinase antibody array

Site specific phosphorylation levels of 43 kinases were measured using the Human Phospho-Kinase Array Kit (catalog no. ARY 003B, R&D Systems) according to the manufacturer’s instructions with the experimental design as described below. Pseudopregnancy was induced in the Wnk1^f/f^ and Wnk1^d/d^ mice as previously described, and mice were euthanized on PPD 4.5. Uterine tissues were frozen at -80°C until ready to proceed. Lysate were extracted independently from 6 mice per group by bead milling in the lysis buffer provided within the kit, and protein concentrations were determined using the BCA Protein Assay Kit (catalog no. 23225, Pierce). Equal amounts from each mouse were pooled in each group (to a total of 900 µg), and the remaining steps followed the standard protocol of the kit. Signal intensity was quantified by ImageJ (59). Images shown in the main figure were chosen to allow visualization of maximal difference between Wnk1^f/f^ and Wnk1^d/d^ mice for selected kinases, but quantification was performed using blots in the non-saturation range.

### Protein extraction from uterine tissues and protein expression analysis

Tissues were homogenized in RIPA Lysis and Extraction Buffer (Thermo Fisher) supplemented with protease inhibitor cocktail (cOmplete Mini, EDTA-free, catalog no. 11836170001, Roche Diagnostics) and phosphatase inhibitor cocktail (phosSTOP, catalog no. 4906837001, Roche Diagnostics), followed by centrifugation at 10,000 X G for 10 minutes at 4°C, and the supernatant was moved into fresh eppendorf. Protein concentrations were measured using the BCA Protein Assay Kit (Pierce). Heat denatured protein samples were resolved using 7.5%, 10% or gradient 4-20% Criterion Tris-HCl precast gels (Bio-Rad), followed by transferring using the Trans-Blot Turbo Transfer System (Bio-Rad), as according to the manufacturer’s instructions. PVDF and nitrocellulose membranes were used for target proteins > 200 KDa and < 200 KDa, respectively. After transfer, the membranes were blocked in 5% w/v non-fat milk or BSA prepared in Tris buffered saline with 0.1% Tween-20 (TBST). Membranes were incubated with primary antibody at 4°C with shaking overnight, followed by three 10 minute washes in TBST the next morning. Membranes were proceeded to secondary antibody incubation at room temperature for at least one hour with shaking, and washed another 3 times in TBST. Depending on the expected signal strength, different peroxidase chemiluminescent substrates were used: KPL LumiGLO^R^ (catalog no. 546101, Seracare), Clarity Western ECL Substrate (catalog no. 1705060, Bio-Rad), and Amersham ECL Prime Western Blotting Detection Reagent (catalog no. RPN2232, GE Healthcare Life Sciences). Antibody sources and dilutions are summarized in table S6. For each western blot, GAPDH or β-tubulin were detected as the loading control, and in cases where the target protein is in the same region as the loading control proteins, a duplicate gel was ran and transferred in parallel. For each set of samples, a representing GAPDH or β-tubulin blot is shown. Details of antibodies used in this study are provided in table S6.

### Tissue clearing and three-dimensional reconstruction

Uterine tissues were fixed in 4% PFA for 16 hours, followed by 3 rinses in PBS. Tissues were then incubated in hydrogel monomer solution AP40 (4% v/v acrylamide and 0.25% w/v VA-044 in PBS) for 72 hours at 4°C, protected from light. Oxygen was then removed from the samples using a chamber connected to vacuum and nitrogen, followed by incubation at 37°C for 3 hours to initiate tissue-hydrogel hybridization. Hydrogel was removed from the tissues via 3 PBS washes, and tissues were subsequently incubated in 8% SDS prepared in PBS for 7 days at 37°C with shaking, and the SDS solution replaced twice during incubation. The tissues were then washed 5 times one hour in PBS and blocked in 5% NDS prepared in PBS/triton X-100 with 0.01% of sodium azide. The samples were then incubated in primary antibody prepared in antibody diluent (2% v/v NDS, 0.01% w/v sodium azide in PBST) for 6 days at room temperature with constant rotation, followed by 5 one hour washes in 0.1% v/v Triton in PBS (PBS-T). Secondary antibody was similarly prepared in antibody diluent and incubated for another 6 days at room temperature with constant rotation and protected from light, replacing antibody half way through incubation. Finally, the samples were washed an additional 5 times one hour in PBS-T and incubated in Refractive Index Matching Solution (80% w/v Histodenz (catalog no. D2158, Sigma-Aldrich) prepared in 0.02M phosphate buffer, pH7.5 with 0.1% Tween-20 and 0.01% sodium azide, refractive index = 1.46) for 1-3 days, and samples were mounted in fresh Reflective Index Mounting Solution using a 1 mm deep iSpacer (www.sunjinlabs.com). Details of antibodies used in this study are provided in table S6.

### Cell culture

Human endometrial epithelial cell line HEC1A and telomerase-transformed human endometrial stromal cells (THESC) were obtained from American Type Culture Collection (ATCC, Rockville, MD, USA). HEC1A cells were cultured in McCoy’s 5A modified medium (catalog no. 16600082, Gibco) and the THSEC cells were maintained in DMEM/F12 (1:1) (catalog no. 11330-032, Gibco), both supplemented with 10% fetal bovine serum (FBS, catalog no. 10437-028, Gibco) and 100 U/mL penicillin and 100 µg/mL streptomycin, unless otherwise stated.

### siRNA transfection and drug treatments

Cells were transfected with siRNAs using the Lipofectamine RNAiMax transfection reagent (catalog no. 13778150, Thermo Fisher) following the manufacturer’s protocol. Cells were transfected with 24 – 72 nM siRNA in transfection medium supplemented with 2% charcoal-stripped FBS (catalog no. 12676-029, Gibco) for 24-48 hours before replacing with fresh growth medium. Proteins were harvested from cells 72 hours after transfection unless otherwise stated. The siRNAs used in this study were: nontargeting siRNA (siCTRL, catalog no. D-001810-10-20, Dharmacon), *Wnk1* targeting siRNA (siWNK1, catalog no. L-005362-02-0005, Dharmacon), *Mtor* targeting siRNA (simTOR, catalog no. L-003008-00-0005, Dharmacon), and *Ppp2r1a* targeting siRNA (siPPP2R1A, catalog no. L-060647-00-0005, Dharmacon). AKT and mTOR inhibitors GDC0941 and rapamycin (catalog no. S1065 and S1039, respectively, Selleckchem) were dissolved in DMSO, and cells were treated with 5 µM GDC 0941 and 10 – 40 µM rapamycin for 24 hours, while the control cells received equivalent volumes of DMSO.

### Immunofluorescence of cultured cells

Cells were cultured in 4-chambered coverglass (catalog no. 155382, Thermo Fisher) as described above. Following transfection and/or drug treatment for the appropriate time period, cells were rinsed in cold PBS, fixed in 4% PFA and permeabilized in 0.5% Triton X-100/PBS for 10 and 5 minutes, respectively, at room temperature. Cells were then incubated in blocking buffer (5% v/v NDS, 0.2% v/v fish gelatin (catalog no. G7765, Sigma-Aldrich), 0.2% v/v Tween-20 in PBS) for 30 minutes at 37°C. Primary antibody was diluted in blocking buffer and added to the cells for 60 minutes, followed by secondary antibody for another 60 minutes; both incubation steps were performed at 37°C in a humidified chamber. Finally, cells were rinsed 3 times with 0.2% Tween-20/PBS and coverslipped using a DAPI containing mounting medium (Vectorshield Hardset™ Antifade Mounting Medium, catalog no. H-1400, Vector Laboratories). Details of antibodies used in this study are provided in table S6.

### WNK1 Immunoprecipitation Mass-spectrometry

HEC1A cells were grown to 70% confluency, followed by collection using trypsin. Cells were washed 2 X in cold PBS, followed by resuspension in cell lysis buffer (50 mM Tris-HCl pH 7.5, 150 mM NaCl, 1 mM EDTA, 1% NP-40, 1% sodium deoxycholate, 0.1% SDS, with protease and phosphatase inhibitors added fresh to 1 X). Cells were incubated on ice for 10 minutes, followed by sonication on medium power (3 X 5 seconds). Lysate was centrifuged at 13,000 X G for 10 minutes at 4°C. WNK1 targeting antibody was added at 1:100 to the supernatant, and incubated with rotation at 4°C overnight. Prewashed beads (50% protein A and 50% protein G, catalog numbers 10002D and 10004D, respectively, Thermo Fisher) were added to the immunocomplex and incubated for 30 minutes at room temperature with rotation. Beads were pelleted using a magnetic separation rack, followed by 3 washes in lysis buffer. Beads were heated to 100°C with 3 X SDS buffer (150 mM Tris-HCl pH 6.8, 6% SDS, 0.3% BPB, 30% glycerol, 3% B-mercaptoethanol) for 5 minutes, before electrophoresis through a 7.5% Criterion Tris-HCl precast gel (Bio-Rad). Gel regions were excised from the SDS-PAGE gel and minced, and digests were performed with a ProGest robotic digester (Genomic Solutions) where the gel pieces were destained by incubation in 25 mM ammonium bicarbonate with 50% acetonitrile (v/v) twice for a total of 30 minutes. The gel pieces were dehydrated in acetonitrile, followed by drying under a nitrogen stream, and further incubated with 250 ng trypsin (Promega) for 8 hours at 37°C. The digests were collected, and peptides were re-extracted three times. The extractions were pooled for each sample, lyophilized and resuspended in 20 µL 0.1% formic acid. The protein digests were analyzed by LC/MS on a Q Exactive Plus mass spectrometer (Thermo Fisher) interfaced with a nanoAcquity UPLC system (Waters Corporation), and equipped with a 75 µm x 150 mm BEH dC18 column (1.8 µm particle, Waters Corporation) and a C18 trapping column (18 µm x 20 mm) with a 5 µm particle size at a flow rate of 400 nL/min. The trapping column was positioned in-line of the analytical column and upstream of a micro-tee union which was used both as a vent for trapping and as a liquid junction. Trapping was performed using the initial solvent composition. A volumn of 5 µL of digested sample was injected into the column, and peptides were eluted by using a linear gradient from 99% solvent A (0.1% formic acid in water (v/v)) and 1% solvent B (0.1%formic acid in acetonitrile (v/v)), to 40% solvent B over 60 minutes. For the mass spectrometry, a data dependent acquisition method was employed with an exclusion time of 15 seconds and an exclusion of +1 charge states. The mass spectrometer was equipped with a NanoFlex source and was used in the positive ion mode. Instrument parameters were as follows: sheath gas, 0; auxiliary gas, 0; sweep gas, 0; spray voltage, 2.7 kV; capillary temperature, 275°C; S-lens, 60; scan range (m/z) of 200 to 2000; 2 m/z isolation window; resolution: 70,000; automated gain control (AGC), 2 X 10^5^ ions; and a maximum IT of 200 ms. Mass calibration was performed before data acquisition using the Pierce LTQ Velos Positive Ion Calibration mixture (Thermo Fisher). Peak lists were generated from the LC/MS data using Mascot Distiller (Matrix Science) and the resulting peak lists were searched using the Spectrum Mill software package (Agilent) against the SwissProt database. Searches were performed using trypsin specificity and allowed for one missed cleavage and variable methionine oxidation. Mass tolerance were 20 ppm for MS scans and 50 ppm for MSMS scans.

### Generation of mammalian YFP-WNK1 expression constructs

The coding region of the WNK1 sequence (NM_014823.3) with attL sites and N-terminal TEV cleavage site was synthesized by GeneWiz Inc. and cloned into pUC57 (Kanamycin) plasmid. Gateway Cloning using LR Clonase II mix (Thermo Fisher) was used to transfer the WNK1 sequence into the Vivid Colors pcDNA6.2/N-YFP vectors (Thermo Fisher), which created the mammalian expression vectors with YFP fused to the N-terminal end of WNK1 (Fig. S7, c4161).

### Co-Immunoprecipitation

HEC1A cells were transfected with cYFP or c4161 for 48 hours, followed by trypsinization, 3 washes and resuspension in lysis buffer (50 mM Tris pH8.0, 400 mM NaCl, 0.1% NP-40 and 0.5 mM DTT, with protease and phosphatase inhibitors freshly added to 1 X). The lysate was incubated at 4°C with rotation for 30 minutes. Lysates were centrifuged at 21,100 X G for 10 minutes at 4°C, and the supernatant was added to 1.5 volumes of 25% glycerol, followed by centrifugation at 21,100 X G for 10 minutes at 4°C. Anti-GFP resin slurry was added to the supernatant and nutated for 1 hour at 4°C. Beads were centrifuged at 1,000 X G for 5 minutes, 4°C, followed by 6 washes in 100 µL of PBST in Bio-Spin columns (catalog number 7326204, Bio-Rad). The bound immunocomplexes were eluted via 0.1 M glycine, pH 2.0, and eluent was neutralized using 2M Tris-HCl, pH 8.0.

### Confocal Microscopy

All fluorescent images presented in this study were captured using the Zeiss LSM 780 UV confocal microscope.

### Statistics

GraphPad Prism versions 7 and 8 were used for data analysis. Each set of data points were first subjected for normality tests. Student’s *t* tests and Mann-Whiteney tests were performed for normally distributed data and non-normally distributed data, respectively. For % of mice with implantation post mating, Fisher’s exact test was performed. In each case, a *p-*value less than 0.05 was considered as significant.

## Supporting information

Supplemental Figures_Chi2020

## Author Contributions

Conceptualization, R.A.C., S.P.W. and F.J.D.; Methodology, R.A.C. and F.J.D.; Validation, R.A.C.; Formal Analysis, R.A.C. and T.W.; Investigation, R.A.C.; Resources, S.L.Y., J.L., C.L.H. and F.J.D.; Data Curation, R.A.C. and T.W.; Writing – Original Draft, R.A.C.; Writing – Review & Editing, R.A.C. and F.J.D.;Visualization, R.A.C.; Supervision, F.J.D.; Project Administration, R.A.C. and F.J.D.; Funding Acquisition, F.J.D.

## Acknowledgements

This work was supported in part by Intramural Research Program of the National Institute of Health (Z1AES103311-01 (F.J.D.)); the Eunice Kennedy Shriver National Institute of Child Health & Human Development (RO1 HD042311 (J.P.L)); and National Institute of Diabetes and Digestive and Kidney Diseases (RO1 DK111542 (C.-L.H)). The authors thank Dr Sheng Song for guidance on the CLARITY technique; Dr Nyssa Adams for conducting the initial breeding trial, Dr Carmen Williams and Dr Sophia Tsai for reviewing the manuscript. We appreciate support from the NIEHS animal facility, Knockout Mouse Core, Digital Imaging Core, the Epigenomics and DNA Sequencing Core, the Fluorescent Microscopy and Imaging Core, the Mass Spectrometry Research and Support Group, the Structural Biology Core of NIEHS for their support and guidance with specialized techniques, as well as the Ligand Assay and Analysis Core at the University of Virginia.

## Supplemental Information Titles and Legends

**Figure S1. Generation of uterine WNK1 ablation mouse model.** (A) The Wnk1^f/f^ mice harbours 2 loxP sites (red block “L”) flanking the second exon of the *Wnk1* gene, which were crossed to mice carrying Cre under the control of PGR promoter. Resultant PGR^Cre/+^; Wnk1^f/f^ mice would have exon2 excised from the genome in PGR expressing cells. (B and C) Decreased *Wnk1* gene and WNK1 protein expression were confirmed using qPCR (B) and western blotting (C) of uterine RNA or protein, respoctively, results shown are mean ± SD, * *p* < 0.05.

**Figure S2. Delayed implantation in the Wnk1**^**f/f**^ **mice was not due to aberrant ovarian function.** (A) Ovulation and fertilization was examined by inducing super-ovulation in the Wnk1^f/f^ and Wnk1^d/d^ mice, followed by mating to wild-type male mice. Number of 2-cell embryos were quantified by oviductal flushing and counting under brightfield microscope. (B and C) Serum progesterone (P4) and estradiol (E2) on GD 4.5 in Wnk1^f/f^ and Wnk1^d/d^ mice. Results shown are mean ± SD.

**Figure S3. Principle component analysis of uterine transcriptome (RNA-seq) during receptivity from 4 Wnk1**^**f/f**^ **mice (red) and 5 Wnk1**^**d/d**^ **mice (blue).** Each dot represent uterine tissues from one mouse, and was clustered based on the transcriptomic profie.

**Figure S4. Quantification and validation of the phosphokinase array.** (A) The phosphokinase array was subjected to densitometrical quantification using ImageJ, and phosphokinases with phosphorylation change (Wnk1^d/d^ density/Wnk^f/f^ density) which exceeded 1.5 fold are shown. (B) Phosphorylation and total levels of AKT, GSK-3α/β, and PRAS40 in the Wnk1^f/f^ and Wnk1^d/d^ uteri on PPD 4.5 were assayed by Western blot, with each lane representing one mouse.

**Figure S5. mTOR is activated in the Wnk1**^**d/d**^ **uteri, but does not regulate AKT activity.** (A) Rapamycin, an mTOR inhibitor was unable to reverse WNK1 ablation induced FOXO1 nuclear exclusion, indicating that AKT activity is maintained. (B) Rapamycin treatment did not rescue WNK1 ablation induced phosphorylation of AKT and FOXO1 in both HEC1A (left) and THESC (right) cells. (C) Expression and phosphorylation of FOXO1 and AKT in HEC1A cells transfected with siCTRL, siWNK1, simTOR, or both siWNKT and simTOR. Results showed no rescue of FOXO1 and AKT phosphorylation by mTOR knock-down.

**Figure S6. WNK1 immunoprecipitation confirmation.** HEC1A cells were subjected to immunoprecipitation (IP) using IgG (negative control) or WNK1 targeting antibody. The pulldown lysate showed enrichment of WNK1 in the WNK1 IP, but not in the IgG (black arrow indicates expected position of WNK1), as determined by western blotting.

**Fig. S7. Plasmid map of the WNK1 mammalian expression construct pcDNA-6.2-N-YFP-WNK1 (c4161).**

**Table S1. Serum progesterone levels in Wnk1**^**f/f**^ **and Wnk1**^**d/d**^ **mice on PPD 4.5 to confirm pseudopregnancy.**

**Table S2. List of differentially expressed genes in the Wnk1**^**d/d**^ **uteri compared to Wnk1**^**f/f**^ **uteri on PPD 4.5.**

**Table S3. Functional annotation of altered transcriptome in Wnk1**^**d/d**^ **uteri at PPD 4.5 (DAVID).**

**Table S4. List of upstream regulators with altered activites in Wnk1**^**d/d**^ **uteri at PPD 4.5 (IPA).**

**Table S5. List of potential WNK1 interacting partners identified by WNK1 IP-MS.**

**Table S6. List of antibodies, sources, and experimental conditions used in this study.**

