## Supplemental Figures_Chi2020 for "WNK1 regulates uterine homeostasis and its ability to support pregnancy"

### Slide 1
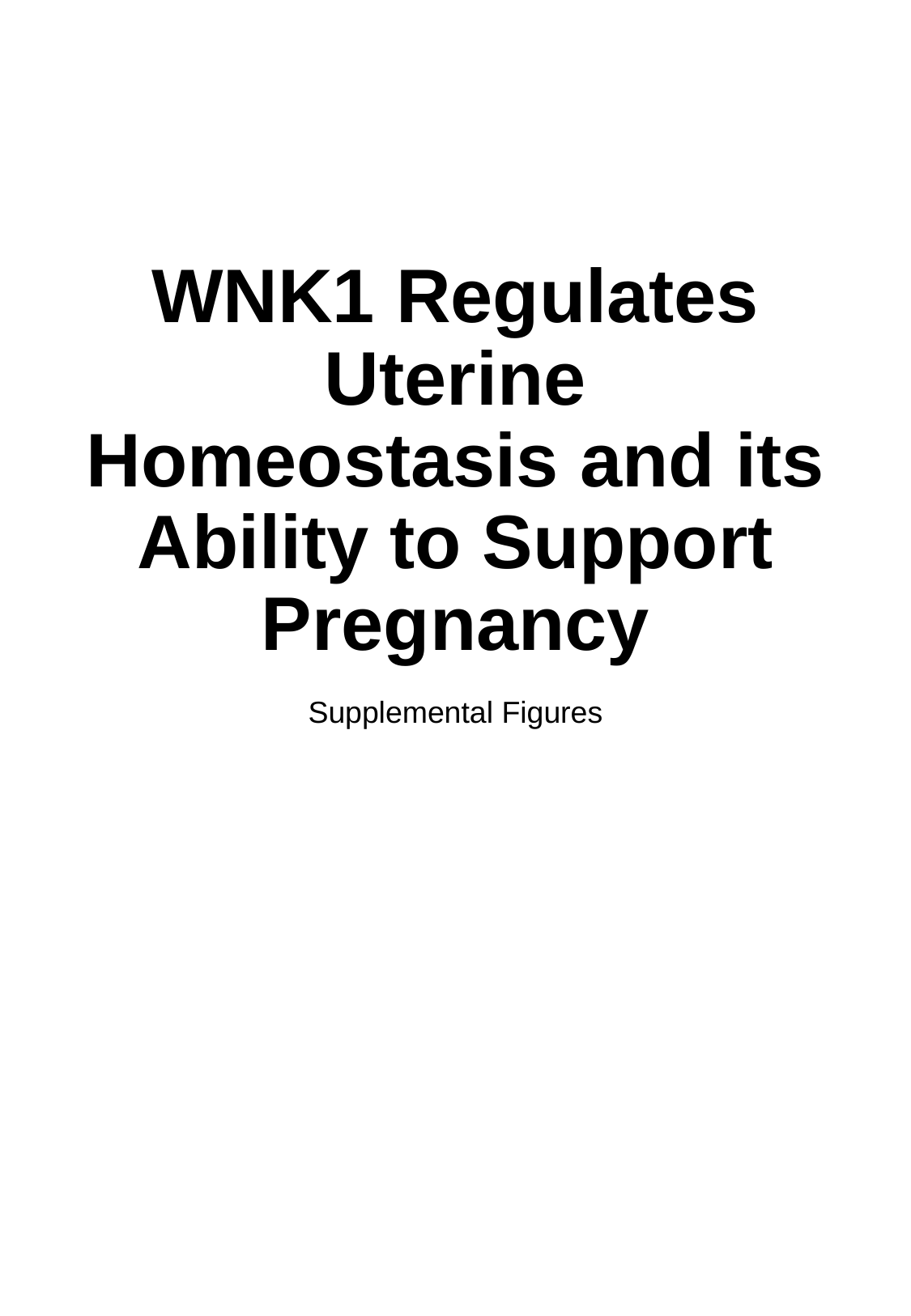

#### WNK1 Regulates Uterine Homeostasis and its Ability to Support Pregnancy
Supplemental Figures

### Slide 2
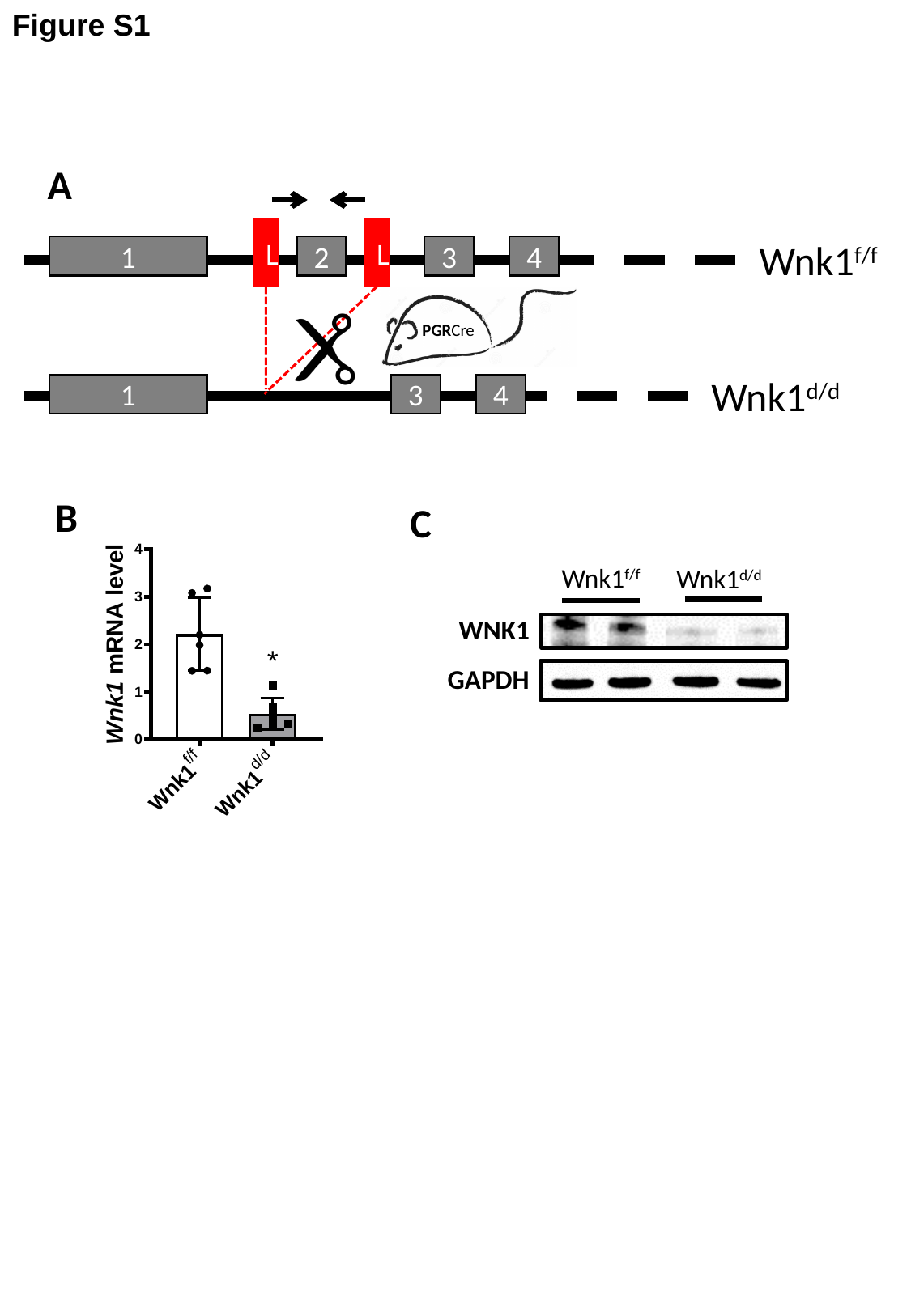

Figure S1
A
L
L
Wnk1^f/f^
4
3
2
1
PGRCre
Wnk1^d/d^
4
1
3
B
C
Wnk1^f/f^
Wnk1^d/d^
WNK1
GAPDH

### Slide 3
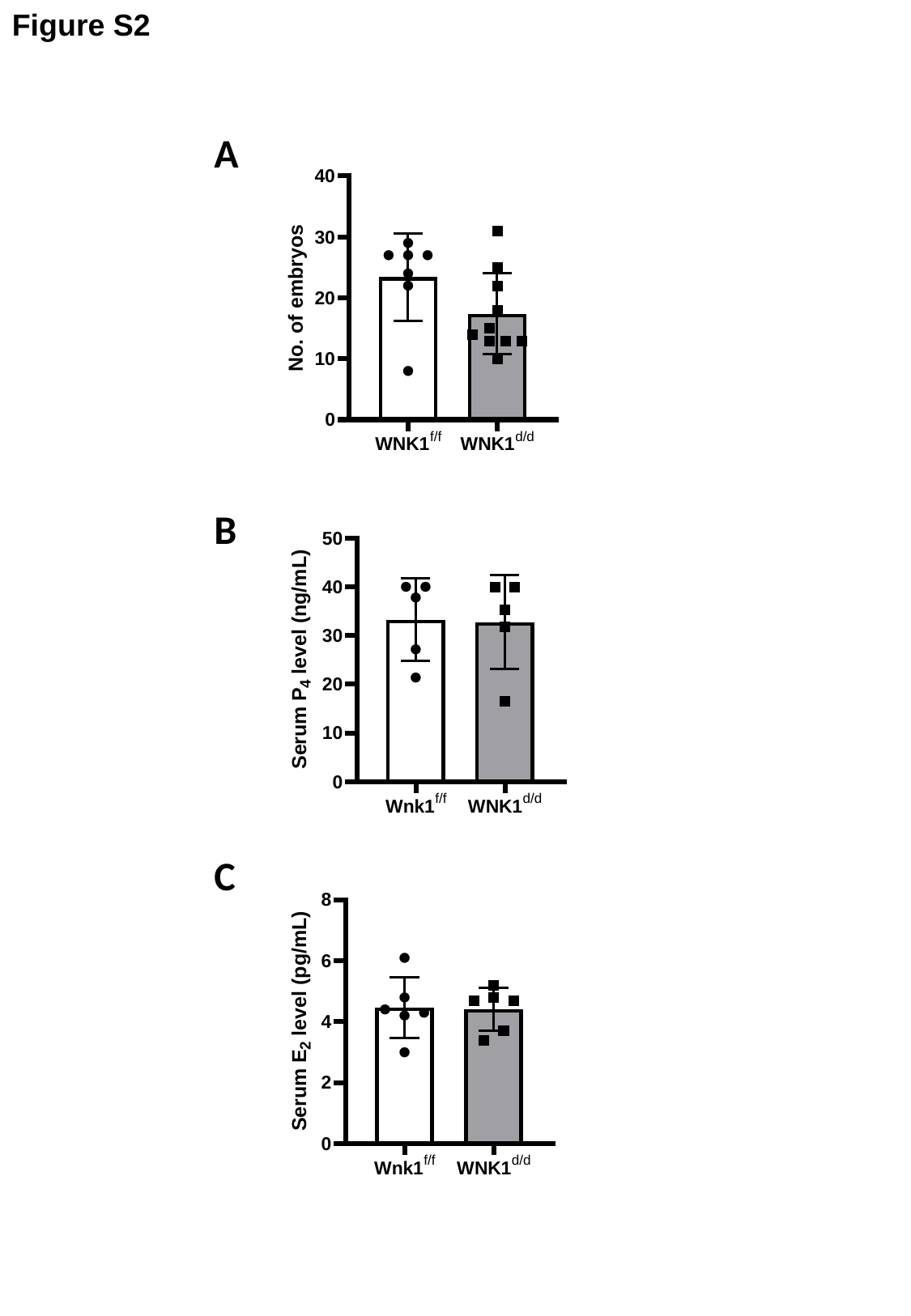

Figure S2
A
B
C

### Slide 4
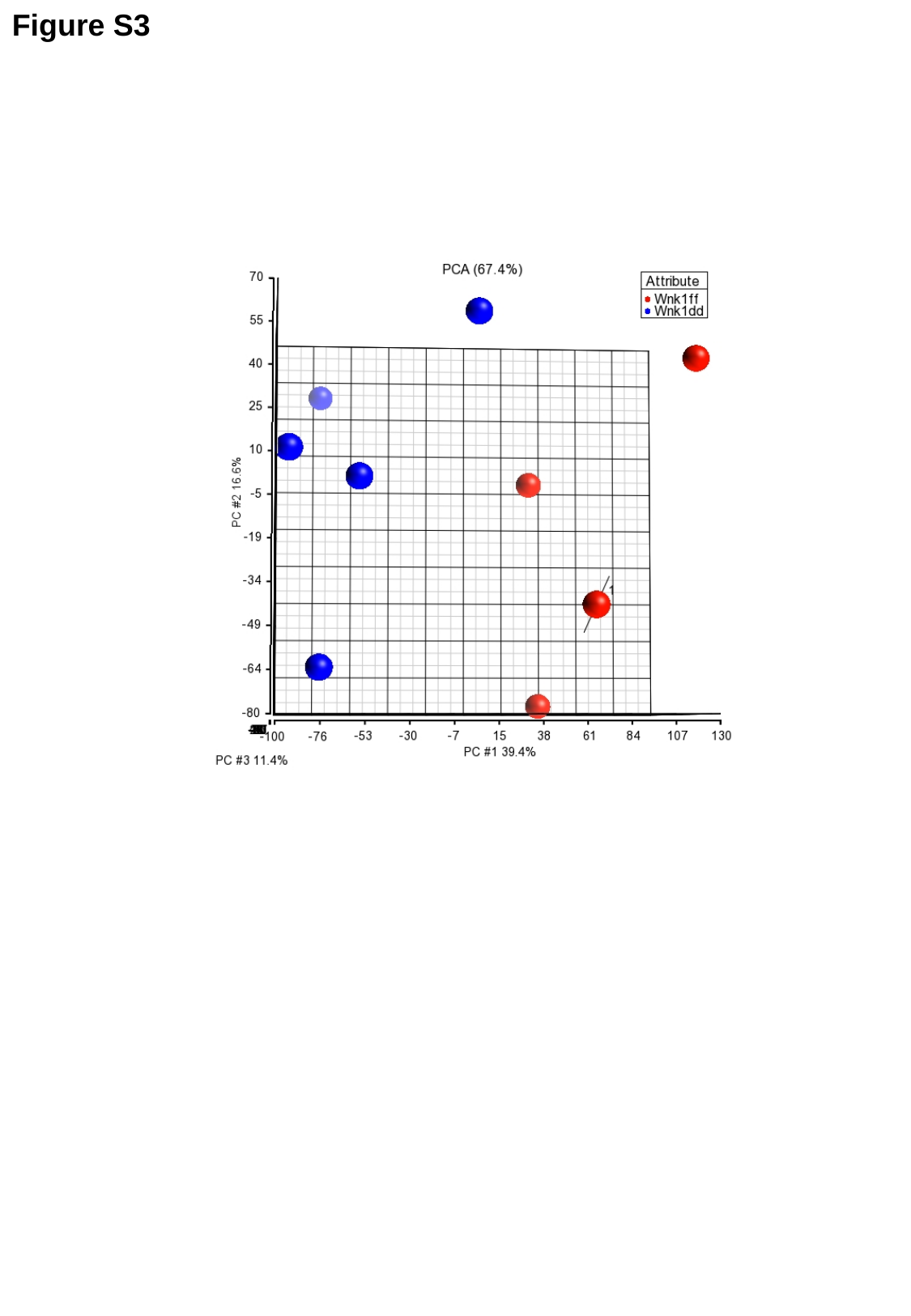

Figure S3

### Slide 5
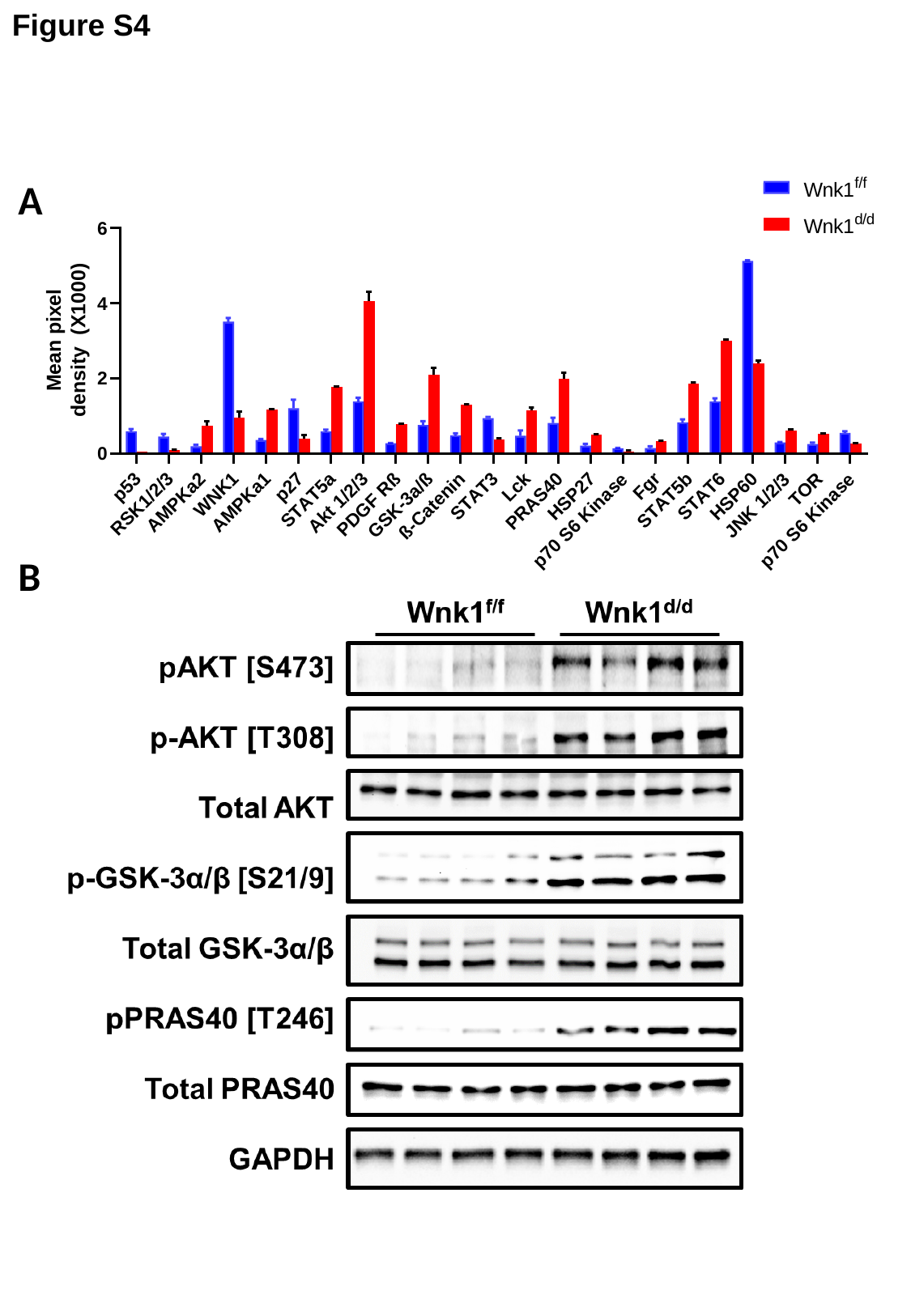

Figure S4
A
B

### Slide 6
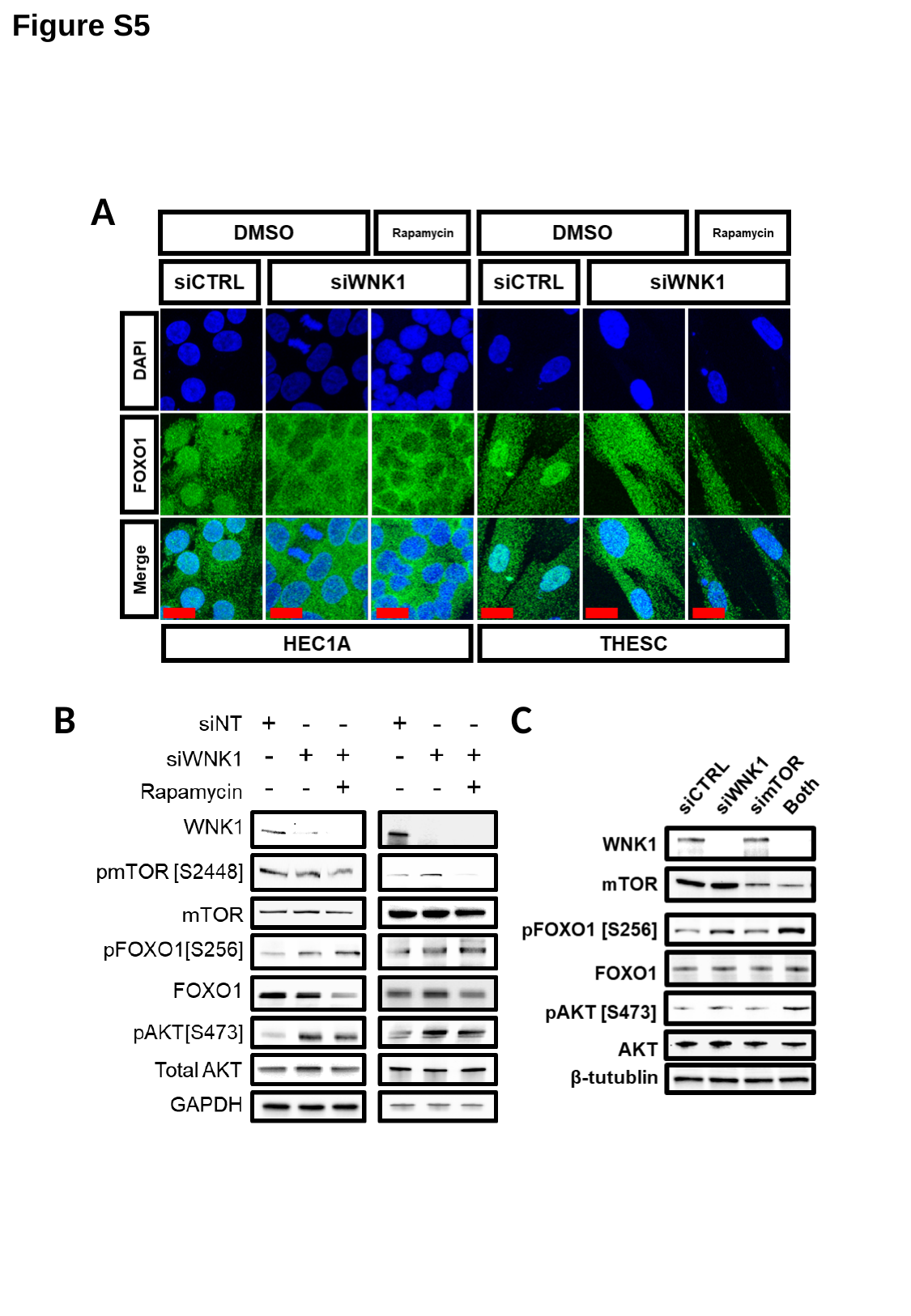

Figure S5
A
B
C

### Slide 7
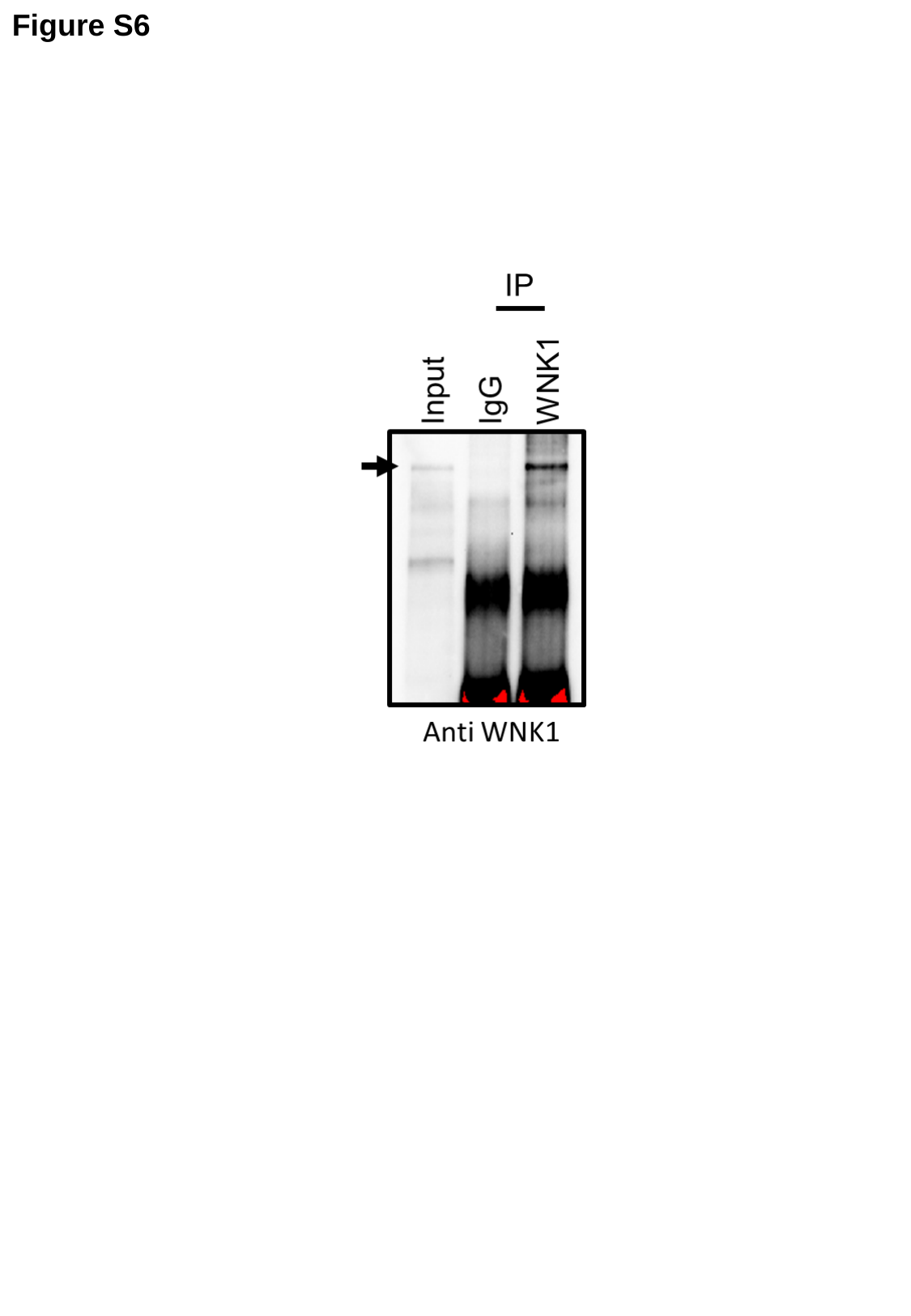

Figure S6

### Slide 8
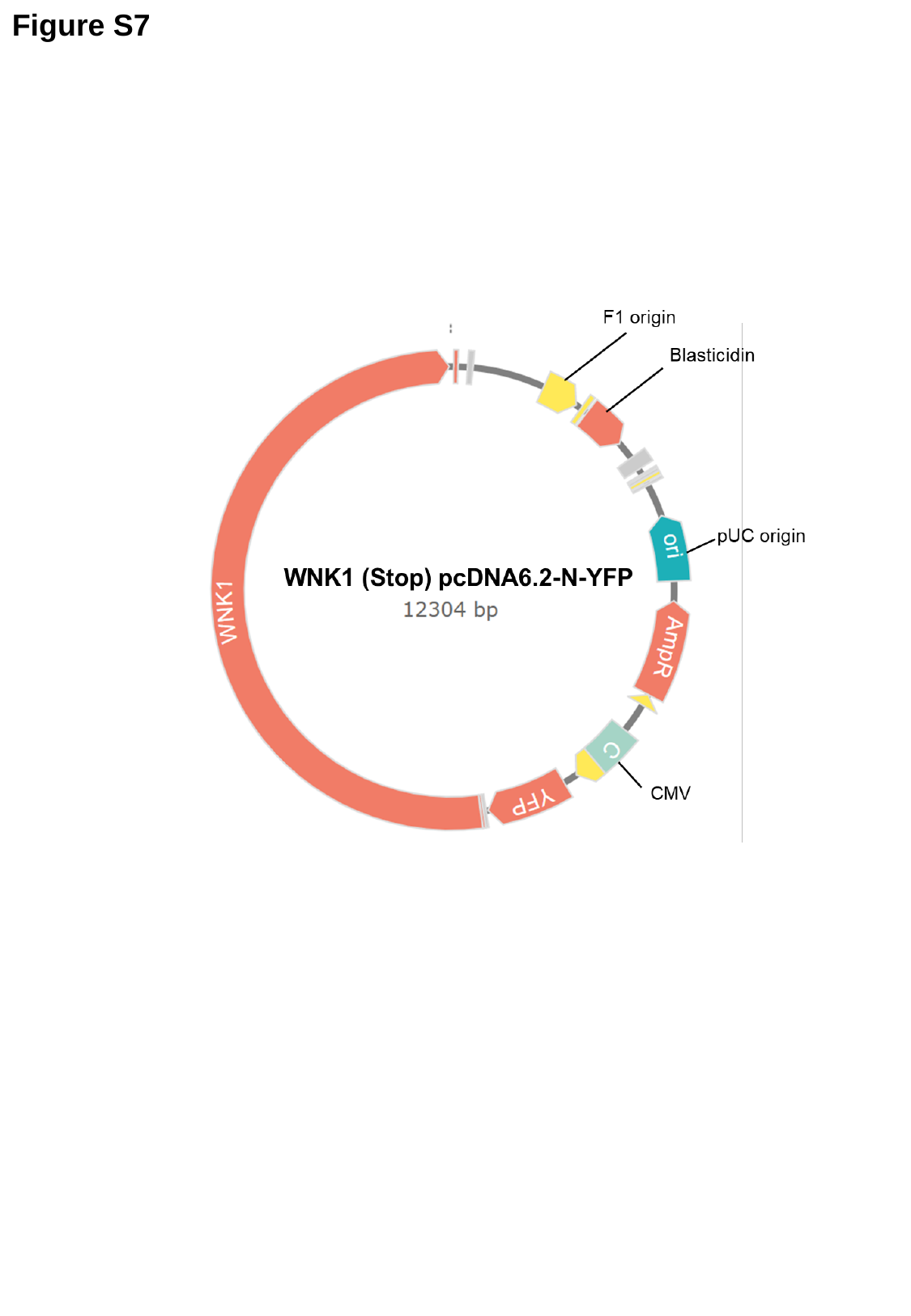

Figure S7
